# AbaM Regulates Quorum Sensing, Biofilm Formation and Virulence in *Acinetobacter baumannii*

**DOI:** 10.1101/2020.11.17.387936

**Authors:** Mario López-Martín, Jean-Frédéric Dubern, Morgan R. Alexander, Paul Williams

## Abstract

*Acinetobacter baumannii* possesses a single divergent *luxR/luxI*-type quorum sensing (QS) locus named *abaR/abaI*. This locus also contains a third gene located between *abaR* and *abaI* which we term *abaM* that codes for an uncharacterized member of the RsaM protein family known to regulate *N*-acylhomoserine lactone (AHL) dependent QS in other β- and γ-proteobacteria. Here we show that disruption of *abaM* via a T26 insertion in *A. baumannii* strain AB5075 resulted in increased production of N-(3-hydroxydodecanoyl)-L-homoserine lactone (OHC12) and enhanced surface motility and biofilm formation. In contrast to the wild type and *abaI*::T26 mutant, the virulence of the *abaM*::T26 mutant was completely attenuated in a Galleria mellonella infection model. Transcriptomic analysis of the *abaM*::T26 mutant revealed that *abaM* differentially regulates at least 76 genes including the *csu* pilus operon and the acinetin 505 lipopeptide biosynthetic operon, that are involved in surface adherence, biofilm formation and virulence. A comparison of the wild type, *abaM*::T26 and *abaI*::T26 transcriptomes, indicates that *abaM* regulates ~21% of the QS regulon including the csu operon. Moreover, the QS genes (*abaI*/*abaR*) were among the most upregulated in the *abaM*::T26 mutant. *A. baumannii lux*-based *abaM* reporter gene fusions revealed that *abaM* expression is positively regulated by QS but negatively auto-regulated. Overall, the data presented in this work demonstrates that *abaM* plays a central role in regulating *A. baumannii* QS, virulence, surface motility and biofilm formation.

**IMPORTANCE:** *Acinetobacter baumanni* is a multi-antibiotic resistant pathogen of global healthcare importance. Understanding *Acinetobacter* virulence gene regulation could aid the development of novel anti-infective strategies. In *A. baumannii*, the *abaR* and *abaI* genes that code for the receptor and synthase components of an *N*-acylhomoserine (AHL) lactone-dependent quorum sensing system (QS) are separated by *abaM*. Here we show that although mutation of *abaM* increased AHL production, surface motility and biofilm development, it resulted in the attenuation of virulence. *abaM* was found to control both QS-dependent and QS-independent genes. The significance of this work lies in the identification of *abaM*, an RsaM ortholog known to control virulence in plant pathogens, as a modulator of virulence in a human pathogen.

## INTRODUCTION

*Acinetobacter baumannii* is a Gram-negative opportunistic nosocomial pathogen that causes a wide range of infections in humans, most commonly pneumonia, but also bacteremia, skin, soft tissue and urinary tract infections, meningitis and endocarditis (1). The rise of multi-drug resistant strains has limited the treatment options for this pathogen which has become a major threat to hospital patients worldwide (2). Indeed, the WHO classified *A. baumannii* as a critical pathogen for which new antibiotics are urgently required (3). For this reason, a better understanding of the virulence of *A. baumannii* should aid the development of new therapeutic strategies for preventing and treating *Acinetobacter* infections. Several virulence factors and regulators involved in *A. baumannii* pathogenesis have been characterized to date. These include outer membrane proteins (e.g. OmpA), pili, capsular polysaccharide, iron acquisition systems, outer membrane vesicles, secretion systems and phospholipases (4–9), as well as regulators such as H-NS and two component systems (10–13). Some *A. baumannii* strains also undergo phase variation where opaque colony variants exhibit greater motility and virulence but reduced biofilm formation compared with the translucent variants (11). A detailed review of *A. baumannii* virulence can be found in Morris et al, 2019 (14).

One well-established mechanism of virulence gene regulation in diverse pathogens is quorum sensing (QS) (15). This cell-cell communication system is employed by bacteria to coordinate the expression of specific genes as a function of population density. QS is mediated via the synthesis, release and detection of diffusible signalling molecules such as the N-acyl-homoserine lactones (AHLs) (16). *A. baumannii* and related pathogenic *Acinetobacter* spp. possess a LuxR/LuxI QS system, consisting of an AHL synthase (*abaI*) and a transcriptional regulator (*abaR*) that is activated on binding an AHL. Most pathogenic *Acinetobacter* spp. produce AHLs with acyl side chains of 10 to 12 carbons length with N-(3-hydroxydodecanoyl)-L-homoserine lactone (OHC12) being most commonly encountered. Many strains are however capable of producing other AHLs (17). Several reports have linked QS to biofilm formation and surface motility (18–20) while others have suggested that it plays a role in virulence in a strain and animal model dependent manner (21, 22). However, our current knowledge of the role of QS in the virulence of pathogenic *Acinetobacter* spp. is limited.

Located adjacent to the *abaI* gene in *Acinetobacter* there is an ortholog of the RsaM protein family. These are found in diverse β- and γ-proteobacteria, including *Burkholderia* spp., *Pseudomonas fuscovaginae, Halothiobacillus neapolitanus and Acidithiobacillus ferrooxidans* (23). The first ortholog to be characterized was RsaM in the plant pathogen *P. fuscovaginae*. This was shown to negatively regulate AHL production and was required for full virulence in rice plants (24). Transcriptomic analysis revealed that RsaM partially regulates the QS regulon, as well as modulating the expression of diverse genes in a QS-independent manner (25). Similarly, TofM, the RsaM ortholog found in *Burkholderia glumae*, represses AHL production while positively regulating toxoflavin and virulence in rice (26). The two RsaM orthologs present in *Burkholderia thailandensis* are both negatively auto-regulated while being positively controlled by their cognate QS systems (27). In *Burkholderia cenocepacia* H111, *Bc*RsaM downregulates AHL biosynthesis and modulates swarming motility, biofilm formation, protease and siderophore production (28). Structural and biochemical analysis of BcRsaM showed that it forms dimers in solution and does not appear to bind DNA or AHLs, suggesting that RsaM family proteins act as post-transcriptional or post-translational regulators (23).

RsaM orthologs clearly play a central role in the regulation of QS-dependent and QS-independent gene expression and virulence in plant pathogens. Here we investigated the role and regulation of *abaM* in *A. baumannii* AB5075, a comparatively recently isolated multi-antibiotic resistant, hypervirulent strain (29). We show that *abaM*, in the opaque variant of A. baumanni 5075, controls AHL production, surface motility and biofilm formation and is required for virulence in a *Galleria mellonella* infection model. QS positively regulates *abaM* expression which is turn is negatively autoregulated. Transcriptomic analysis of the *abaM* and *abaI* mutants indicate that the *abaM* and QS regulons overlap. These data are consistent with a central role for *abaM* in regulating gene expression and the pathobiology of *A. baumannii*.

## RESULTS

### Organization of the QS locus in *A. baumannii* 5075

The genome of *A. baumannii* AB5075 possesses a single QS locus comprised of two divergently transcribed genes: an AHL synthase (*abaI/ABUW_3776/ABUW_RS18385*) and a response regulator (*abaR/ABUW_3774/ ABUW_RS18375*) gene. Between *abaR* and *abaI* a third gene is located which here we term *abaM* (*ABUW_3775*) (**Fig. 1A**). The chromosomal organization of these three QS genes is well conserved among *Acinetobacter* spp (**Fig. S1**). Despite the location of *abaM* adjacent to *abaI* and transcribed in the same direction, *abaM* has only low (~20-30 %) sequence identity to orthologs present in *Pseudomonas fuscovaginae* and *Burkholderia spp* (**Fig. 1B**). However, it retains the well-conserved regions shared by other RsaM orthologues, including most of the protein secondary structural elements and the characteristic hydrophobic core cluster, consisting of four tryptophan residues (Trp60, Trp75, Trp77 and Trp125) (23).

**FIG 1.**
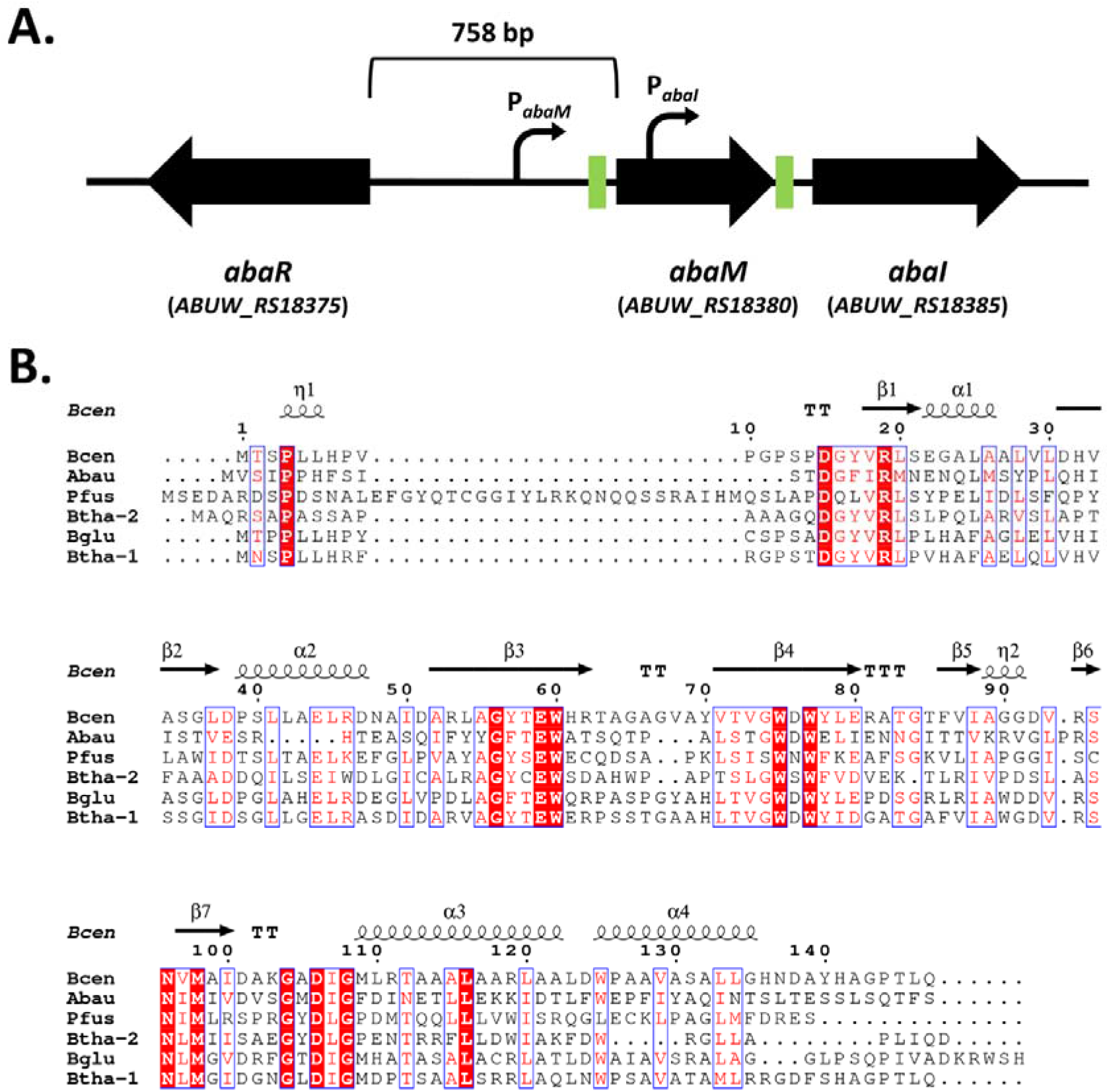
**(A)** Schematic of the *abaR*MI quorum sensing locus in *A. baumannii* AB5075 showing the organization of the three QS genes and their orientation. Green boxes represent predicted *lux* boxes. Curved arrows represent the predicted *abaM* and *abaI* promoters. **(B)** Multiple sequence alignment of *A. baumannii* AB5075 *abaM* (Abau) with previously characterized orthologs in other bacterial species: Bcen: *Burkholderia cenocepacia* J2315 *Bc*rsaM, Pfus: *Pseudomonas fuscovaginae* UPB0736 *rsaM*, Btha-2: *Burkholderia thailandensis* E264 *rsaM*-2, Bglu: *Burkholderia glumae* BGR1 TofM, *Burkholderia thailandensis* E264 *rsaM*-1. The MUSCLE algorithm (47) was used to create the alignment and ESPript (48) to render residue similarities and generate the final figure. White characters in a red background indicate conserved residues. Red residues indicate conservative substitutions. Blue frames indicate highly conserved regions. The secondary structures in *B. cenocepacia Bc*rsaM (PDB entry 4O2H) is displayed above the alignment. η: 3_10_-helix, α: α-helices, β: β-strands, **TT**: strict β-turns, **TTT**: strict α-turns.

### AHL production is enhanced in an *abaM* mutant

AHL production in the opaque variants of the AB5075 wild type, *abaM*::T26 and *abaI*::T26 mutants respectively was quantified via LC MS/MS during growth under static conditions since it appears to be enhanced by surface attachment in other *A. baumannii* strains (30). OHC12 (**Fig. 2A**) was the major AHL produced by AB5075 under these conditions. Compared with the wild type, the *abaM*::T26 mutant produced significantly greater amounts of OHC12 at each time point (between 100 and 875-fold difference) sampled (**Fig. 2B**). N-(3-hydroxydecanoyl)-L-homoserine (OHC10), was also detected at much lower concentrations in the *abaM*::T26 mutant throughout growth, but only in the 24 h sample in the wild type (**Fig. 2C**). No AHLs were detected in any of the *abaI*::T26 samples (**Fig. 2B and C**). These data suggest that *abaM* is a negative regulator of AHL production.

**FIG 2.**
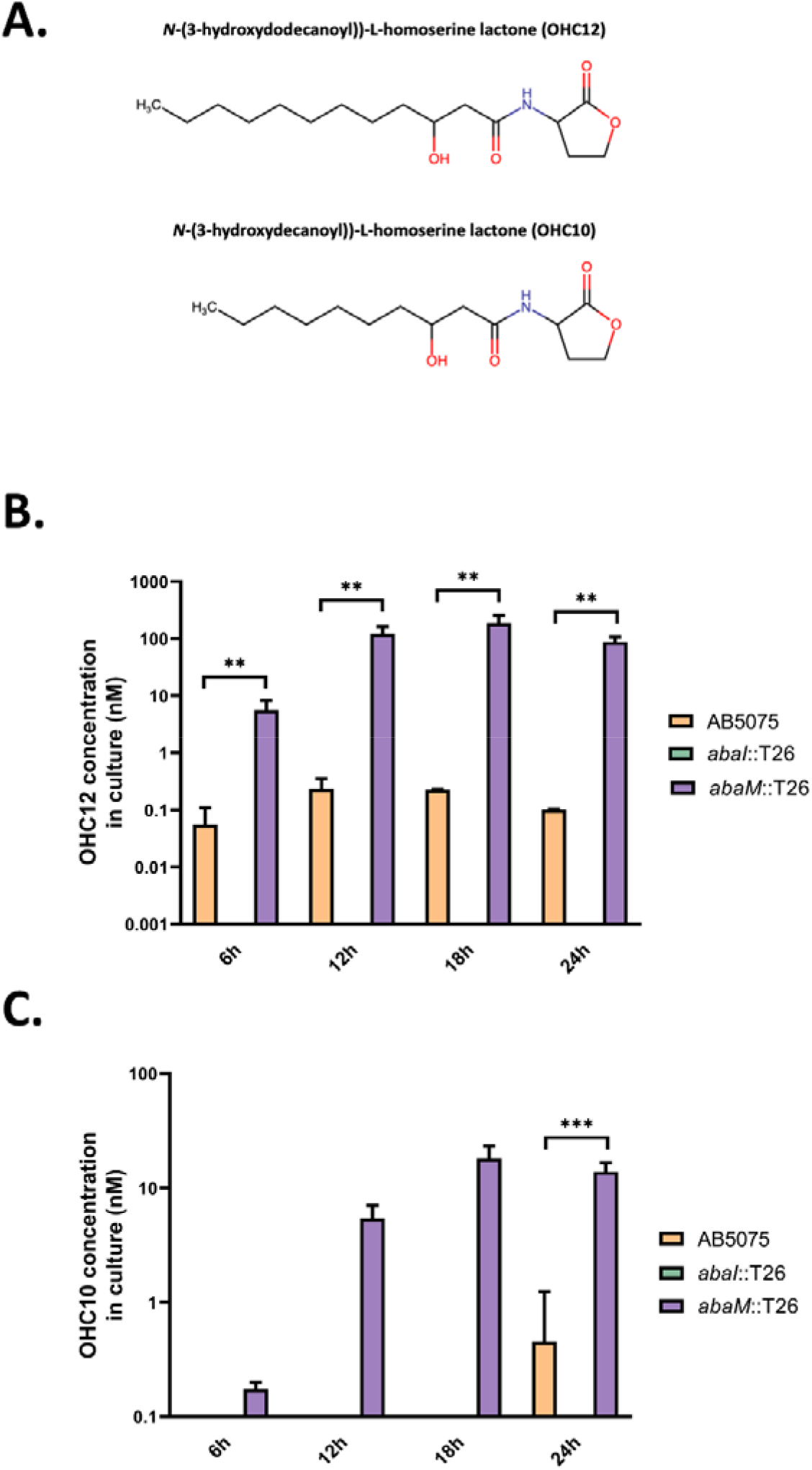
AHL production in wild-type, *abaI*::T26 and *abaM*::T26. **(A)** Chemical structures of the AHLs produced by *A. baumannii* AB5075. Quantification of OHC12 **(B)** and OHC10 **(C)** production throughout growth. Error bars represent the standard deviation between three biological replicates. Asterisks indicate statistically significant differences: **, p ≤ 0.01; ***, p ≤ 0.001

### Contribution of QS and *abaM* to surface motility, biofilm formation and virulence

The surface motility of all three strains on 0.3% Eiken agar LS-LB plates was examined. Compared with the wild-type (59.6 ± 0.7 mm), the *abaI*::T26 mutant exhibited significantly reduced (36.5 ± 1.4 mm) whereas the *abaM*::T26 mutant was significantly more motile (76.7 ± 2.4 mm) (**Fig. 3A**). The provision of exogenous OHC12 increased the surface motility of both wild type and *abaI* mutant to a similar level to that of the *abaM* mutant (**Fig. S2A**)

**FIG 3.**
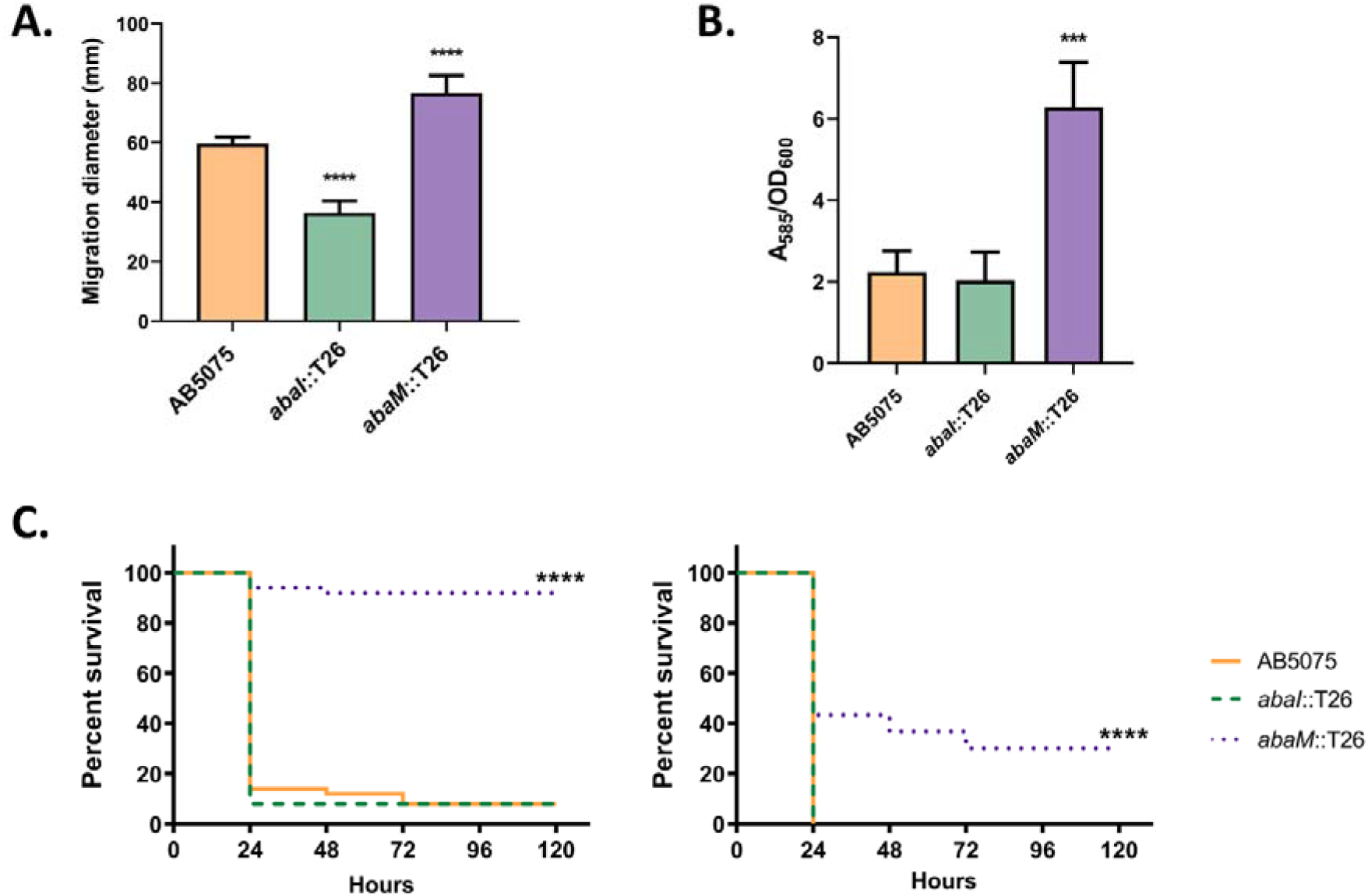
Phenotypic characterization of the *Acinetobacter abaM* and *abaI* mutants. **(A)** Surface motility on agar **(B)** Biofilm formation on polypropylene. For both biofilm and surface motility assays, error bars indicate standard deviation. Asterisks indicate statistically significant differences compared with the wild-type AB5075 strain: ***, p ≤ 0.001; ****, p ≤ 0.0001. **(C)** *Galleria mellonella* larvae killing after inoculation of approximately 2 × 10^4^ (left) or 2 × 10^5^ (right) CFU/larva. Each graph represents data from 3 independent biological replicates together. At least 30 larvae were used for each strain and assay. None of the control larvae died after 5 days. Asterisks indicate statistically significant differences compared with the wild-type AB5075 strain: ****, p ≤ 0.0001.

The ability of AB5075, *abaI*::T26 and *abaM*::T26 to attach to abiotic surfaces was evaluated on propylene tubes. The *abaM* mutant formed ~3 fold more biofilm than the wild-type (**Fig. 3B**). Under these growth conditions, the biofilm produced by the *abaI* mutant (opaque variant; **Fig. 3B**) was not significantly different from than the wild type but increased following the exogenous provision of OHC12 (**Fig. S2B**).

The contribution of *abaM* and QS to AB7075 virulence was assessed using a G. mellonella larvae infection model (**Fig. 3C**). No differences in killing were observed between the wild-type and the *abaI*::T26 mutant when injecting either 2 × 10^4^ or 2 × 10^5^ CFU/larva. However, the *abaM*::T26 mutant was significantly less virulent than the parental strain. Larvae injected with 2 × 10^5^ CFU of the *abaM*::T26 mutant also showed a lower death rate than the larvae injected with 2 × 10^4^ CFU of the wild-type or the *abaI*::T26 mutant. Exogenously supplied OHC12 did not affect the virulence of the wild type (**Fig. S2C**).

Overall, these results suggest that *abaM* is a negative regulator of surface motility and biofilm formation and required for full virulence in *G. mellonella*.

### Genetic complementation of the *abaM* mutant phenotypes

Complementation of the *abaM*::T26 mutant with the *abaM* gene in trans (pMQ_*abaM*) restored surface motility (**Fig.S3A**), biofilm formation (**Fig. S3B**) and reduced both OHC12 and OHC10 production by approximately 50% (**Fig. S3C and D**). However, complementation of the *abaM* mutation did not restore *abaM*::T26 virulence to wild-type (**Fig. S3E**).

### Transcriptomic analysis of *abaI*::T26 and *abaM*::T26

To characterize the *abaM* and QS regulon of *A. baumannii* AB5075, we performed transcriptomic profiling of AB5075 in comparison with the *abaM*::T26 and *abaI*::T26 mutants using RNA sequencing (RNA-seq), which was then validated for two key target genes via quantitative real-time PCR. For these analyses we used total RNA extractions from cells grown for 18 h in static conditions when maximum OHC12 levels are produced by the *abaM* mutant (**Fig. 2**).

When compared with the wild-type strain, 88 genes were upregulated and 9 downregulated in the *abaI*::T26 mutant, while 52 were upregulated and 24 downregulated in the *abaM*::T26 mutant (log2(fold change) ≥ 1; **Fig.4** and **Tables S3** and **S4**). Moreover, 21 of the upregulated genes were shared between *abaI*::T26 and *abaM*::T26 (**Fig. 4, Tables S3** and **S4**), while none of the downregulated genes were co-regulated. Among the genes upregulated in both mutants there were all the genes of the csu operon, a putative TetR family transcriptional regulator (*ABUW_1486/ABUW_RS07245*) located immediately upstream of the csu operon as well as genes coding for a flavohemoprotein, an uncharacterized transcriptional regulator, a thermonuclease, a sulfate permease, a toxic anion resistance protein and 7 hypothetical proteins (**Tables S3 and S4**).

**FIG 4.**
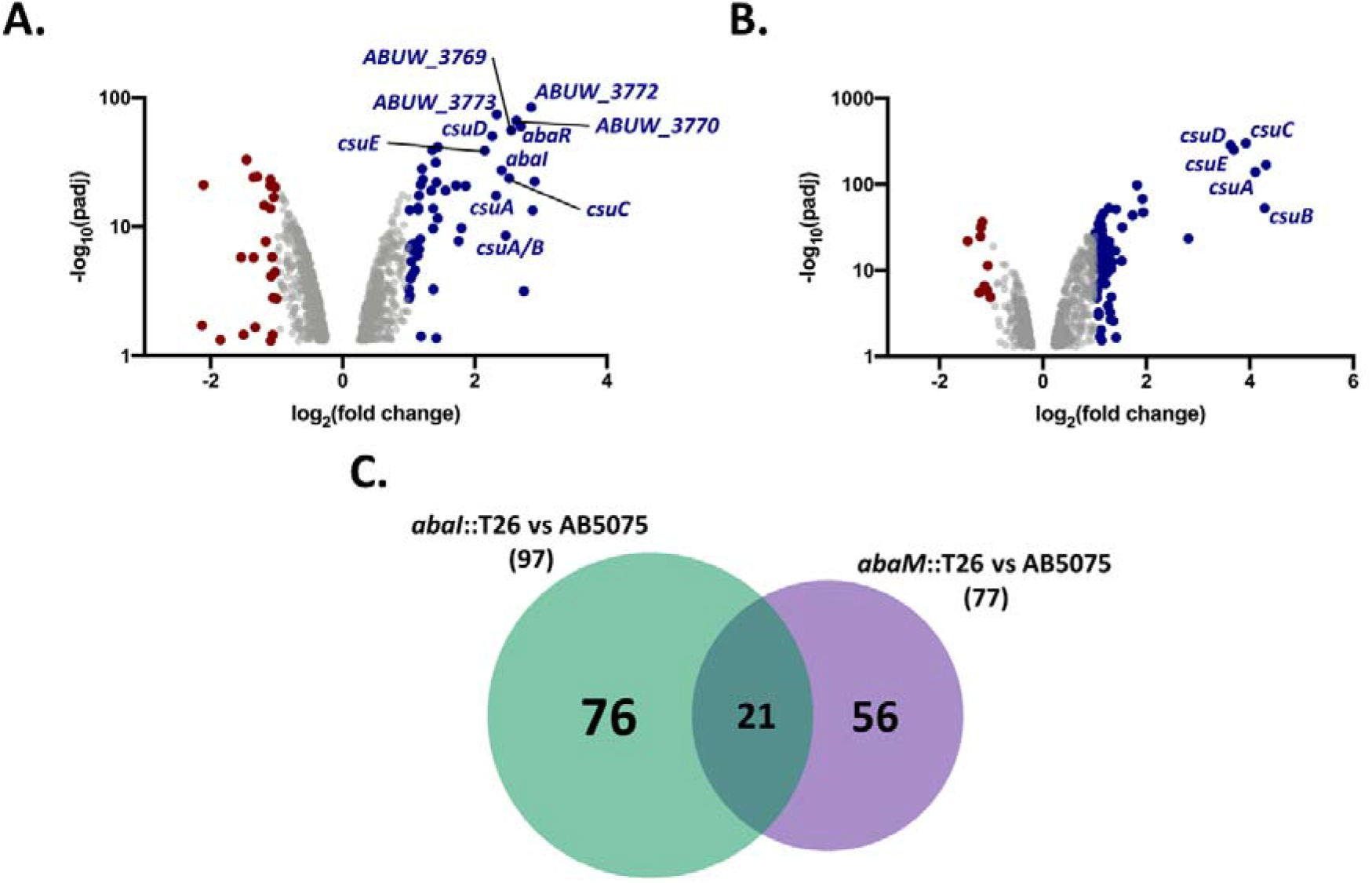
Comparison of the transcriptomes of the *abaI*::T26 and *abaM*::T26 mutants. Genes differentially expressed in **(A)** *abaM*::T26 **(B)** *abaI*::T26 compared with wild-type. Blue circles indicate upregulated genes. Red circles indicate downregulated genes. Grey circles represent genes where changes in expression are unlikely to be biologically significant. **(C)** Venn diagram showing that *abaM* regulates genes that are both QS-dependent and QS-independent.

Moreover, genes of the biosynthetic operon involved in the synthesis of acinetin 505, the QS response regulator, *abaR* and the AHL synthase *abaI* were both upregulated in the *abaM*::T26 mutant, which also differentially expressed genes encoding for proteins involved in the stress response, iron-acquisition, diverse metabolism and energy production, chaperones, protein folding and antibiotic resistance e.g. class D beta-lactamase OXA-23 involved in resistance to carbapenems (**Table S4**). Similarly, differentially regulated genes in the *abaI* mutant included diverse metabolic and energy production-related genes, as well as diverse genes coding for transcriptional regulators, stress response-related proteins and membrane transport proteins (**Tables S3**).

To validate the transcriptomic data, qPCR was performed with the same RNA samples used in the RNA-seq for the *csuA/B* and the *ABUW_3773* (the first gene of the acinetin 505 biosynthetic operon) genes (**Fig. 5**). When compared with the wild-type, the data obtained showed a significant increase in *csuA/B* expression in both the *abaI*::T26 and *abaM*::T26 mutants, while ABUW_3773 expression was significantly higher only in the *abaM*::T26 mutant. These results correlate with the data obtained from the RNA-seq experiments.

**FIG 5.**
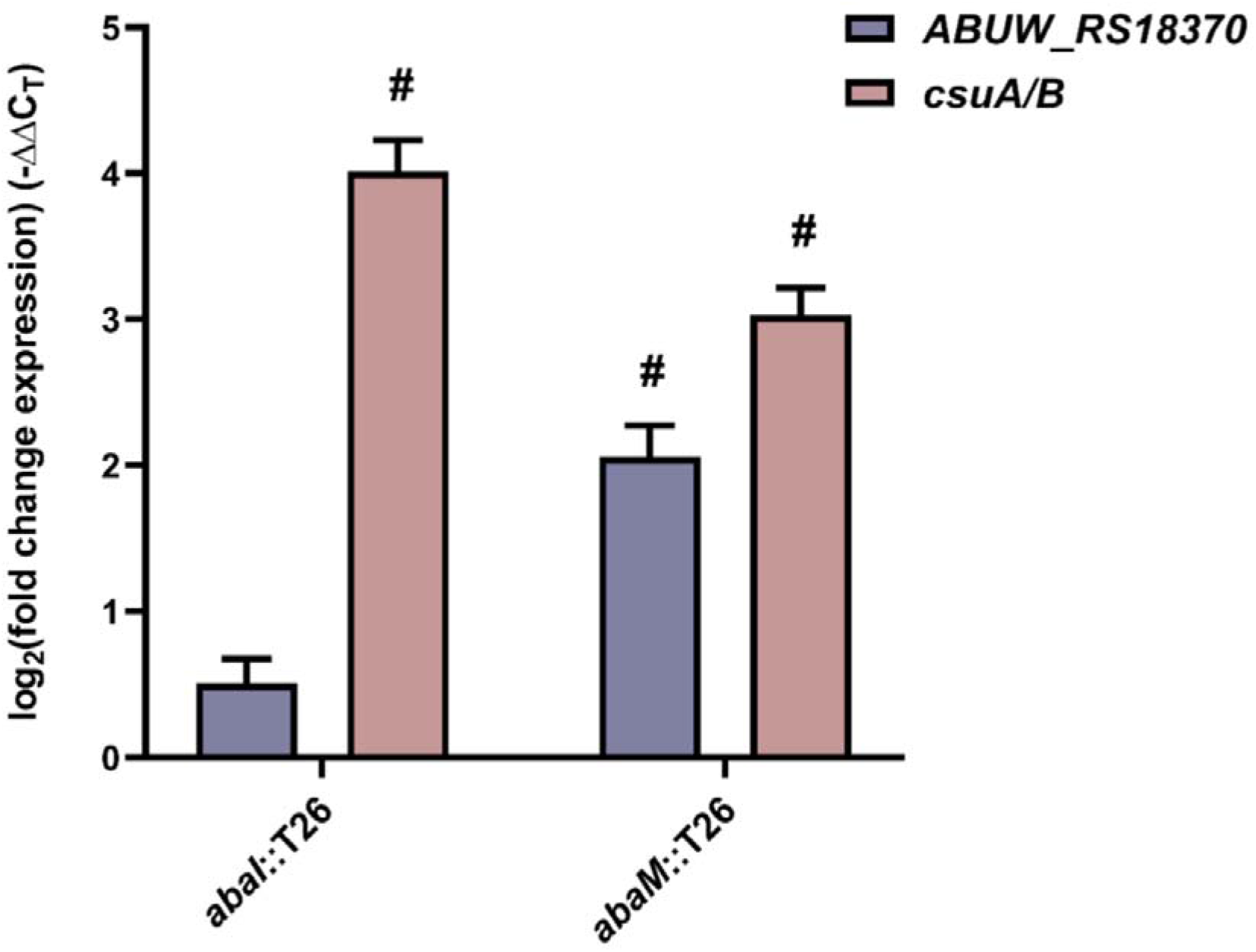
Validation ABUW_3773 and *csu* expression by quantitative real-time PCR. Relative expression of *csuA/B* and *ABUW_3773* in mutants compared with the wild-type AB5075 strain. The expression was normalized in relation to an endogenous control gene (*rpoB*). Error bars indicate standard deviation between 3 independent biological replicates. Hashtags (#) indicate a biologically significant difference (|log2(fold change)| ≥ 1) compared to the wild-type.

### Regulation of *abaM*

To further elucidate the regulation of *abaM* expression, an *abaM* promoter – *luxCDABE* operon fusion was constructed. This was introduced via a miniTn7 transposon into AB5075 and both *abaI*::T26 and *abaM*::T26 mutants, and the activity of the predicted promoter was measured by luminescence output in the presence or absence of OHC12. The activity of the *abaM* promoter significantly varied between the strains. In the *abaM*::T26, luminescence was approximately 40% higher than in the wild-type, while the *abaI*::T26 mutant showed a 75% reduction when compared with the parental strain (**Fig. 6A**). Moreover, exogenous provision of OHC12 increased the *abaM* promoter activity in all three strains (**Fig. 6B**). Similarly, **Fig. 6C** shows that expression of an *abaI*::*lux* promoter fusion which is reduced in the *abaI* mutant compared with the wild type strain, is strongly stimulated by OHC12. These data suggest that *abaM* expression is negatively autoregulated but, in common with *abaI* is positively regulated by QS which in turn is negatively controlled via *abaM* (**Fig. 7**).

**FIG 6.**
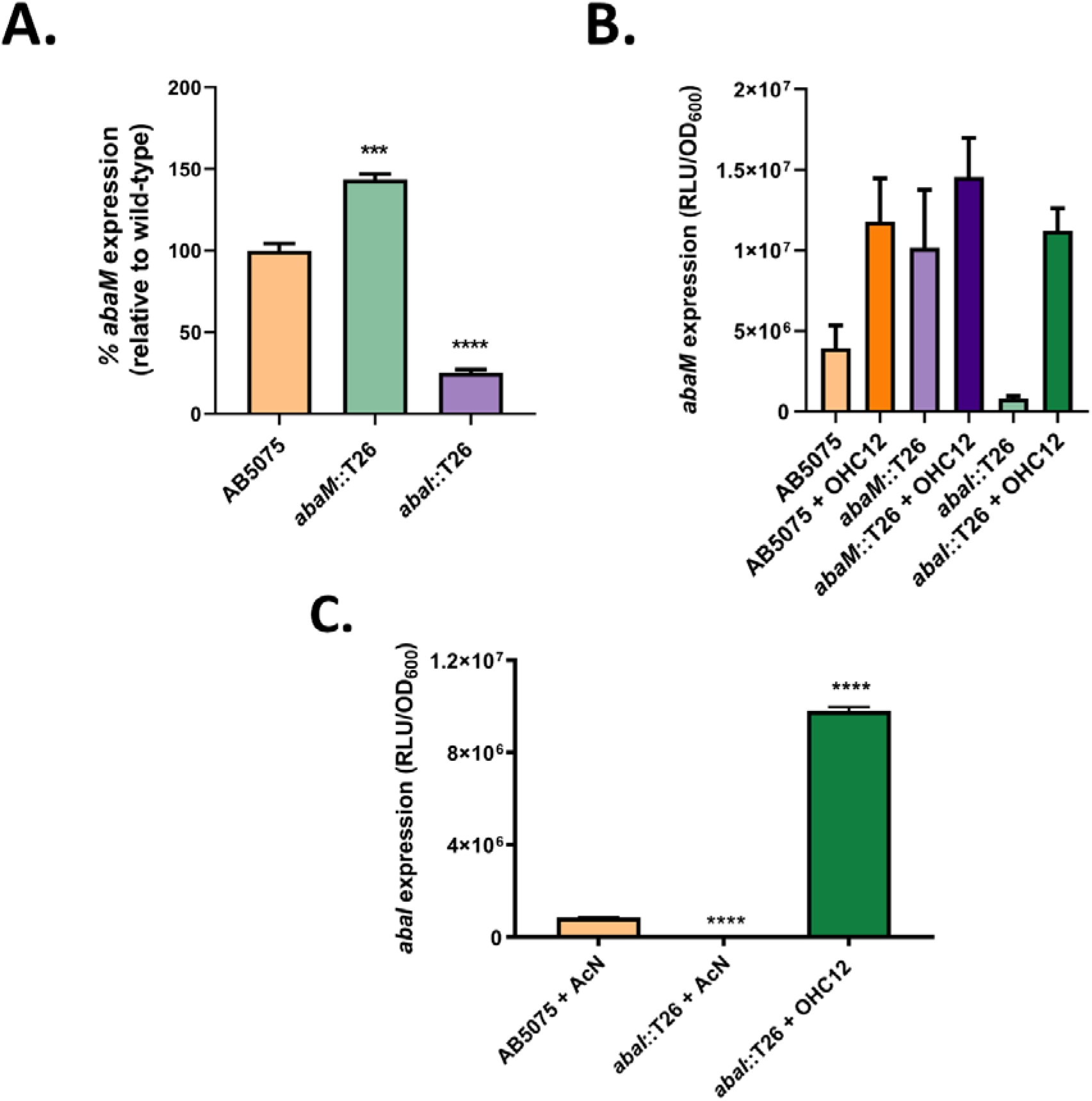
Expression of *abaM* and *abaI*. **(A)** *abaM* promoter activity in *A. baumannii* A5075 wild type, *abaM*::T26 and *abaI*::T26 relative to the wild-type strain. **(B)** *abaM* promoter activity in response to exogenous OHC12 as a function of growth (RLU/OD_600_). **(C)** *abaI* promoter activity in wild type, *abaI* mutant and in response to exogenous OHC12 as a function of growth (RLU/OD_600_). Error bars indicate standard deviation between 3 independent biological replicates. Asterisks indicate statistically significant differences compared to the wild-type AB5075 strain: ***, p ≤ 0.001; ****, p ≤ 0.0001.

**FIG 7.**
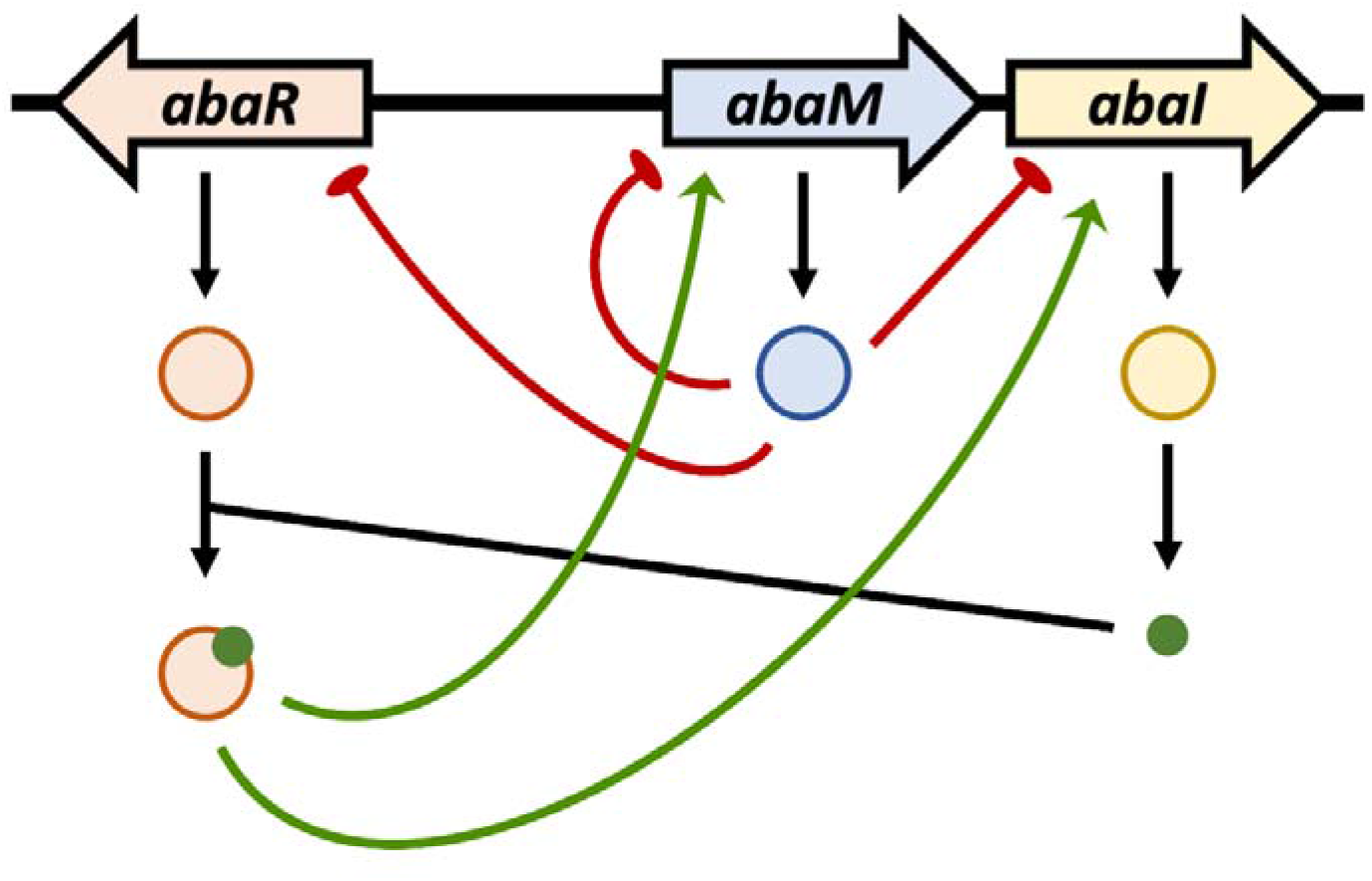
Proposed model for the QS/*abaM* incoherent feed forward loop (IFFL) in *A. baumannii* 5075. *abaR* activated by OHC12 positively activates expression of *abaM* and *abaI* and hence OHC12 production. *abaM* is negatively autoregulated and also represses expression of both *abaR* and *abaI*. Under the growth conditions used here, *abaM* negatively regulates surface motility, biofilm and *Csu* pili but is required for virulence as *abaM* mutants are avirulent in *Galleria mellonella* larvae. Green arrows and red blunt ended lines represent positive and negative regulation respectively

## DISCUSSION

In this study we have established that the RsaM ortholog, *abaM* plays a major role in regulating QS-dependent and QS-independent genes in *A. baumannii* 5075. Disruption of *abaM* substantially increased AHL production, indicating that *abaM* negatively regulates AHL biosynthesis. In *A. baumannii* 5075, the concentration of OHC12 produced was very low (< 1 nM), suggesting that, at least under the static growth conditions used in this study, *abaM* exerts tight control over QS. This raises the question of when and under what conditions QS is active in AB5075, especially since the half-maximal responses for LuxR proteins activated by long chain AHLs is in the 5-10 nM range (31). In *A. baumannii*, the QS locus has an *RXI* topological arrangement previously described in other bacteria, where the *X* gene between the QS regulator (R) and the synthase (I) genes is a negative regulator of QS. Examples of three different classes of X include *rsaL*, *rsaM* and *mupX* (32). In this context, *abaM* appears to behave similarly to *RsaL* in the Pseudomonas aeruginosa *lasR/*RsaL*/lasI* QS system, despite their functional and structural differences. The *RsaL* gene which is divergent to lasI is positively regulated by LasR and antagonizes LasR-mediated activation of lasI so counteracting the QS positive feedback autoinduction and providing AHL homeostasis (33). This type of regulatory circuit (**Fig. 7**), termed an incoherent feed forward loop (IFFL) in contrast to simple feed forward loops, can display complex behaviours that include stabilization of output signals and bounded output which ensures robustness against fluctuations in the input signal levels (32, 34). Hence the large increase in AHL production following deletion of *abaM*, *RsaL* and *rsaM* genes that result in less virulent mutants is indicative of the importance of a stabilizing negative regulatory pathway in AHL-dependent QS systems (32, 34).

Consistent with this model, *abaM* expression was found to be negatively auto-regulated but positively regulated by QS. *In silico* analysis of the DNA sequence between *abaR* and *abaI* using Bprom (www.softberry.com) and BDGP (www.fruitfly.org) as well as our RNAseq data all predict the presence of putative −10 and −35 regions for *abaI* and *abaM* respectively suggesting that these genes are not co-transcribed. Similar findings have been reported for *rsaM1* and *rsaM2* from *Burkholderia thailandensis* (27). Reverse transcriptase (RT)-PCR (**Fig. S4**) confirm that *abaM* and *abaI* in *A. baumannii* A5075 do not form an operon. Moreover, the *abaM* gene has a predicted lux-box (CTGGTTAAATATAACAG) 68 bp upstream of the start codon and 178 bp downstream of the −35 and −10 promoter elements. This lux-box is similar to those found upstream of *abaI* (CTGTAAATTCTTACAG) in both *A. baumannii* 5075 and *A. nosocomialis* M2. These results are consistent with an IFFL circuit although further work will be required to fully characterize its properties and control of genes co-regulated by QS and AbaM.

Phenotypic characterization of the A. baumanni *abaM* mutant revealed enhanced surface motility and biofilm formation but reduced virulence compared with the wild type. Previous studies have shown that *rsaM* orthologues are required for full virulence in plants (24, 26), but this is, to the best of our knowledge, the first time that an *rsaM*-like gene has been reported to be required for full virulence in a human pathogen albeit in an insect infection model. The contribution of Bc*rsaM* in *B. cenocepacia* to swarming motility and biofilm formation has also been reported (28). However, deletion of bc*rsaM* reduced both swarming and surface attachment, the opposite to that observed for the *abaM* mutant. Interestingly, both biofilm formation and surface motility in *Acinetobacter* have been associated with increased virulence (10, 35).

Genetic complementation of the *abaM* mutant was achieved for surface motility, biofilm formation and AHL production but not for virulence in *Galleria mellonella*. Similar observations have been previously reported for other *abaM* orthologues, most notably tofM (26) and bc*rsaM* (28), leading to the suggestion that *rsaM*-like proteins may be cis-acting regulators (23).

To further define the role of QS in *A. baumannii* 5075, phenotypic characterization of the AHL synthase mutant, *abaI*::T26, was performed. The mutant did not produce any detectable AHLs, consistent with previous studies and bioinformatic analysis indicating that *A. baumannii* 5075 possesses a single QS locus that is responsible for AHL production (18, 36). Similarly, disruption of *abaI* negatively affected surface motility and responded to exogenous 3OHC12, as previously noted for other *Acinetobacter* strains/species (20). In AB5075, biofilm formation in the opaque variant of the *abaI*::T26 mutant was not significantly different from with wild type. However, it increased well above wild type in response to exogenous OHC12. consistent with other work on the *Acinetobacter* AHL synthase (18, 37). Previous studies on QS and biofilm formation in *Acinetobacter* have been performed with strains that were, in contrast to AB5075, either not phase variable or not known to be phase variable. Since the experiments performed in this study were all carried out with the opaque *Acinetobacter* variant, we also investigated biofilm formation by the translucent variant. **Fig S5** shows that the translucent variant produced less biofilm that the wild type. However, biofilm formation increased for both opaque and translucent variants in response to exogenous OHC12. Furthermore, our results suggest that QS does not play an important role in virulence in the *G. mellonella*. While this is not unprecedented (38), the role of *Acinetobacter* QS in virulence is still not well defined, and other studies suggest that QS may play an important role, depending on the strain and infection model used (21, 22). Overall, our data suggest that for strain AB5075, QS is important in surface motility and biofilm formation, but not virulence. However, further work is required to fully elucidate the role of QS in the pathogenesis of *Acinetobacter* infection and any cross-talk with other regulatory networks.

Here we performed RNAseq on *A. baumannii* AB5075 grown in static conditions where AHL production was elevated in order to identify genes regulated via *abaM* and QS and likely to be involved in surface attachment and biofilm formation. A comparison of the genes differentially expressed in the *abaM* and *abaI* mutants with the wildtype revealed that *abaM* has both QS-dependent (~22% of the QS regulon) and QS independent gene targets. A similar overlap has been noted for the *rsaM* regulon and its cognate QS system in *P. fuscovaginae* (25). Among the most upregulated genes in both *abaI* and *abaM* mutants when compared with the wild-type were those belonging to the *csu* operon (*ABUW_1487-ABUW_1492/ABUW_RS07250-ABUW_RS07275*). This operon encodes the proteins responsible for the synthesis of the *Csu* pilus, a type I chaperone-usher pilus involved in attachment and biofilm formation (39–41). Moreover, the *abaM*::T26 mutant also showed higher expression of some genes of the acinetin 505 biosynthetic operon, which has also been linked to biofilm formation in *A. baumannii* ATCC17978 (42).

A previous comparison of the transcriptomes of the multi-drug resistant clinical A. baumanni strain 863 with an isogenic *abaI* deletion mutant highlighted the differential regulation of 352 genes involved in carbon source metabolism, energy production, stress response and translation (49). However, apart from *abaI*, no other common differentially regulated genes could be identified when the A5075 *abaI* and the 863 *abaI* mutant transcriptomes are compared. This may be because of the different strains and growth conditions and sampling times used. In addition, the *abaI* mutant reported by Ng et al (49) exhibited a growth defect (49). This raises the possibility of a secondary mutation contributing to the transcriptome data which was not validated by chemical or genetic complementation with OHC12 or *abaI* respectively.

The work described here establishes that *abaM* plays a central role in regulating QS, surface motility, biofilm formation and virulence. The apparently contradictory regulatory impact of *abaM* and *abaI* mutations that result in either increased or no AHL production respectively on the expression of genes such as the *csu* cluster can be explained as follows. In an *abaI* mutant (no AHLs), *abaM* expression is reduced and hence *csu* expression is increased. In an *abaM* mutant *csu* expression is also increased as *abaM* is absent (**Fig. 7**). Further work will be required to elucidate the biochemical function and mechanism of action of *abaM* and the *rsaM* protein family in general.

## MATERIALS AND METHODS

### Strains and growth conditions

The strains and plasmids used are listed in **Table S1**. *A. baumannii* AB5075 (29) and the isogenic *abaI*::T26 and *abaM*::T26 mutants were obtained from the transposon library available from the University of Washington (44). *A. baumannii* was routinely grown in low-sodium chloride (5g/L) lysogeny broth (LS-LB). OHC12 was synthesized as described before (46). The opaque and translucent phases of the wild type, and mutants were separated as described by Tipton *et al* (11) after growth on phase-observation LB (PO-LB) plates and observation of colonies under light microscopy using oblique indirect illumination.

### Construction of a genetically complemented *abaM*::T26 strain

Plasmid pMQ557M (**Table S1**) was obtained by digesting pMQ557 with *Pml*I (to remove the genes required for yeast replication) and re-ligating the resulting large linear product. The *abaM* gene plus 768 bp from its upstream region (containing the predicted native promoter) were amplified by PCR using primers listed in **Table S2**. The PCR fragments were digested with BamHI and KpnI and ligated in the multiple cloning site (MCS) of both pMQpMQ557M and introduced into *abaM*::T26 by electroporation. The stability of the vector pMQ557M and *abaM* complementing plasmid pMQ_*abaM* in both A5075 and the *abaM* mutant mutant were confirmed by repeated daily subculture and plating out on LB agar with and without hygromycin (125μg/ml) to determine viable counts as cfu/ml (**Fig. S6**).

### Construction of *abaM*::luxCDABE and *abaI*::luxCDABE promoter fusions

The *abaR* gene and the intergenic region between *abaR* and *abaM* (for the *abaM* fusion) or the region between *abaR* and the *abaI* (for the *abaI* fusion) were amplified by PCR and ligated in pGEM®-T Easy using the pGEM®-T Easy Vector System (Promega). The resulting plasmids and the promoterless *luxCDABE* operon were digested with KpnI and BamHI and ligated in order to introduce the *lux* operon downstream of the predicted promoter of *abaM* or *abaI*. These constructs were transferred into the MCS of the miniTn7T in pUC18T-miniTn7T-HygR plasmid (**Table S1**) after digesting with NotI and PstI and ligating the corresponding fragments.

MiniTn7T-based constructs were inserted into *A. baumannii* through four-parental conjugation. Briefly, PBS-washed overnight cultures of the *E. coli* DH5α donor strain (containing pUC18T-mini-Tn7T_Hyg^R^_*abaR*_P*abaM*::*lux* or the pUC18T-mini-Tn7T_Hyg^R^_*abaR*_P*abaI*::*lux*), the *E. coli* DH5α helper strain (containing pUX-B13), the E. coli DH5α mobilizable strain (containing pRK600) and the A. baumanii recipient strain were mixed in a 1:1:1:1 ratio and grown on LB agar prior to counterselection with hygromycin (125 μg/ml for miniTn7 selection) and gentamicin (100 μg/ml).

The miniTn7 transposon (45) carrying the *abaM* promoter-*lux* operon fusion was inserted into *A. baumannii* AB5075 and the isogenic *abaI*::T26 and *abaM*::T26 mutants; while the *abaI* promoter-lux fusion was inserted into AB5075 and the isogenic *abaI*:T26 mutant. Bioluminescence output from the reporter fusions as a function of bacterial growth was quantified using an Infinite 200 PRO (Tecan Diagnostics) plate-reader over 24 h and OD600 and relative light units RLUs were recorded every 30 min. When required OHC12 was added at 200 nM unless otherwise stated.

### Biofilm assays

Strains to be tested were inoculated into 1.5 ml polypropylene microcentrifuge tubes in LS-LB with or without OHC12 and incubated under static conditions at 37°C for 24 h. Biofilms were quantified by staining with 0.25% crystal violet, extracting with ethanol and recording the absorbance at A585.

### Surface motility assays

Surface motility was quantified as previously described (11) on LS-LB plates with or without OHC12 and containing 0.3% Eiken agar. Plates were incubated at 30°C for 16 h.

### AHL extraction and detection

Cell-free supernatants from cultures grown in LS-LB under static conditions at 37°C were sterile-filtered and extracted with acidified ethyl acetate. Extracts were evaporated to dryness and subjected to by liquid chromatography with tandem mass spectrometry (LC-MS/MS) as previously described (46).

### *G. mellonella* killing assays

*G. mellonella* larvae (Trularv™) were obtained from BioSystems Technology Ltd, U.K. Assays were performed as described previously (11). Briefly, *Acinetobacter* (2 × 10^4^ or 2 × 10^5^ cfu) were injected into the larval hemolymph, incubated at 37°C and the larvae monitored for viability. At least 10 larvae were used for each strain and assay.

### Total RNA extraction and RNA-seq

Bacteria were cultured in LS-LB under static conditions at 37°C for 18h. Cells were resuspended in RNAprotect (Qiagen) prior to extracting total RNA using an RNeasy minikit (QiAgen). After treatment with DNA-free (Invitrogen) the absence of DNA contamination was conformed using by PCR and the quality and quantity of the RNA samples was established using a 2100 Bioanalyzer (Agilent). Samples were sent for 150 bp paired-end sequencing via an Illumina platform and bioinformatic analysis, to NovoGene (Hong Kong, China). The data have been deposited in NCBI’s Gene Expression Omnibus (50) and are accessible through GEO Series accession number GSE151925 (https://www.ncbi.nlm.nih.gov/geo/query/acc.cgi?acc=GSE151925; token for reviewers: wxqxkwygphkbliv).

### Quantitative real-time PCR (qPCR)

Complementary DNA (cDNA) synthesis and qPCR were carried out using LunaScript RT Supermix and Luna Unive*RsaL* qPCR Master Mix (New England BioLabs), respectively. The oligonucleotides used for qPCR are listed in **Table S2** and qPCR was carried out in triplicate using a 7500 Real-Time PCR System (Thermofisher). Negative controls lacking template or RNA incubated without reverse transcriptase were included. The housekeeping gene rpoB was used as endogenous control for normalization.

### Reverse transcription PCR (RT_PCR)

cDNA was amplified using Q5 High-Fidelity polymerase (New England Biolabs) using specific primers annealing in the coding region of each gene. Genomic DNA, extracted using DNeasy Blood and Tissue kit (QIAgen), was used as a positive control. The PCR products were run in a 1.5% agarose electrophoresis gel before imaging under UV light using a Gel Doc XR+ Imager (Bio-Rad).

## Supporting information

SuppInformation

## ACKNOWLEDGEMENTS

We thank Nigel M. Halliday for AHL quantification and Alex Truman for AHL synthesis, Robert M.Q. Shanks (University of Pittsburgh) for kindly providing the pMQ557 plasmid and Prof. Philip N. Rather from Emory University for his guidance and advice on *Acinetobacter* phase variation. This work was supported via Wellcome Trust joint senior investigator awards to MRA and PW (grant nos. 103882 and 103884). MLM was supported by a University of Nottingham postgraduate doctoral studentship.

## Notes

### Competing Interest Statement

The authors have declared no competing interest.

### Summary of Updates

The supplemental section has now been added

