## Supplementary material for "AbaM Regulates Quorum Sensing, Biofilm Formation and Virulence in *Acinetobacter baumannii*": SuppInformation

### SUPPLEMENTAL MATERIAL

Table S1: Strains and plasmids used in this study

| Strain or plasmid | Description | Reference or source |
| --- | --- | --- |
| STRAINS | | |
| *Escherichia coli* | | |
| *E. coli* K-12 DH5α | F^–^ *endA1* *glnV44* *thi-1 recA1 relA1 gyrA96 deoR* *nupG purB20* φ80d*lacZ*ΔM15 Δ(*lacZYA*-*argF*)U169 hsdR17(r_K_^–^m_K_^+^) λ^–^ | Lab stock |
| *Acinetobacter baumannii* | | |
| AB5075 | Wild-type parental strain of the hypervirulent, multi-drug resistant *A. baumannii* AB5075, isolated from a patient with tibia osteomyelitis. | (Jacobs *et al*, 2014) |
| *abaM*::T26 | AB5075 with the T26 transposon disrupting the *rsaM* orthologue *abaM* (*ABUW_3775*). | (Gallagher *et al,* 2015) |
| *abaI*::T26 | AB5075 with the T26 transposon disrupting the quorum sensing synthase gene *abaI* (*ABUW_3776*). | (Gallagher *et al*, 2015) |
| AB5075 P*abaM*::*lux* | AB5075 carrying the mini-Tn*7*T_Hyg^R^_P*abaM*::*lux* insertion. | This study |
| AB5075 P*abaI*::*lux* | AB5075 carrying the mini-Tn*7*T_Hyg^R^_P*abaI*::*lux* insertion. | This study |
| *abaM*::T26  P*abaM*::*lux* | *abaM*::T26 carrying the mini-Tn*7*T_Hyg^R^_P*abaM*::*lux* insertion. | This study |
| *abaI*::T26  P*abaM*::*lux* | *abaI*::T26 carrying the mini-Tn*7*T_Hyg^R^_P*abaM*::*lux* insertion. | This study |
| *abaI*::T26  P*abaI*::*lux* | *abaI*::T26 carrying the mini-Tn*7*T_Hyg^R^_P*abaI*::*lux* insertion. | This study |
| AB5075  pMQ557M | AB5075 carrying the pMQ557M plasmid. | This study |
| *abaM*::T26  pMQ557M | *abaM*::T26 carrying the pMQ557M plasmid. | This study. |
| *abaM*::T26  pMQ_*abaM* | *abaM*::T26 carrying the complementation plasmid pMQ_*abaM*. | This study |
| PLASMIDS | | |
| pGEM®-T Easy | Commercial plasmid for PCR cloning. | Promega |
| pBluelux | pBlueScript II KS(+) (Agilent Technologies) plasmid containing the promoterless *luxCDABE* operon. | Lab stock |
| pBluescript-IISK_KO_Multi_HYG_UTRs | pBlueScript II KS(+) (Agilent Technologies) containing the hygromycin resistance cassette (*hph*). | Lab stock |
| pGEM_ *abaR*_P*abaM*::*lux* | pGEM^®^-T Easy containing *abaR* gene, the *abaR*-*abaM* intergenic region and the *abaM* promoter – *lux* operon fusion cloned in the MCS. | This study |
| pGEM_ *abaR*_P*abaI*::*lux* | pGEM^®^-T Easy containing the region from the *abaR* gene to the *abaI* promoter – *lux* operon fusion cloned in the MCS. | This study |
| pUC18T-mini-Tn*7*T | pUC18 plasmid with the mini-Tn*7* containing only terminator regions and a multi-cloning site (MCS). | (Choi *et al*, 2005) |
| pUC18T-miniTn*7*T_Hyg^R^ | pUC18T-mini-Tn*7*T with the *hph* cassette cloned in the MCS. | This study |
| pUC18T-mini-Tn*7*T_Hyg^R^_*abaR*_P*abaM*::*lux* | pUC18T-mini-Tn*7*T_Hyg^R^ with the *abaR* gene, the *abaR*-*abaM* intergenic region and the *abaM* promoter – *lux* operon fusion cloned in the MCS. | This study |
| pUC18T-mini-Tn*7*T_Hyg^R^_*abaR*_P*abaI*::*lux* | pUC18T-mini-Tn*7*T_Hyg^R^ with the region from the *abaR* gene to the *abaI* promoter – *lux* operon fusion cloned in the MCS. |  |
| pUX-B13 | Plasmid carrying the transposase genes required for four-parental conjugation. | Lab stock |
| pRK600 | Helper plasmid carrying the mobilization genes required for four-parental conjugation. | Lab stock |
| pMQ557M | pMQ557 vector, replicative and stable in  *A. baumannii* containing the *hph* cassette as antibiotic marker, with the yeast replication genes removed. | Robert Shanks (University of Pittsburgh) |
| pMQ_*abaM* | pMQ557M containing the *abaM* gene and 758 bp of the upstream region. | This study. |

Table S2: Primers used in this study. Bases corresponding to the restriction sites are underlined.

| Primer | Sequence (5’ 🡪 3’) | | Description |
| --- | --- | --- | --- |
| P*abaM*::*lux*_FW | CTCCGATTAATTATAATTAACC | | Construction of the P*abaM*::*lux* and P*abaI*::*lux* reporters |
| P*abaM*::*lux*_RV | GCGCGGATCCGCGCGGTACCCCATGCTACCTGCTTAAGTACCC | | Construction of the P*abaM*::*lux* reporter |
| P*abaI*::*lux*_RV | GGATCCGGTACCCATTACAAGTGCTTCCACTTA | | Construction of the P*abaI*::*lux* reporter |
| *rpoB*_qPCR_FW | TCGTGTTGAGCGTGCTGTTA | qPCR of the endogenous control gene *rpoB* | |
| *rpoB*_qPCR_RV | TGCAGCAGCAACTGGTTTTG | qPCR of the endogenous control gene *rpoB* | |
| *csuAB*_qPCR_FW | AGCAGCAACAGGTGGCAATA | qPCR of the *csuA/B* gene | |
| *csuAB*_qPCR_RV | GGTCTGTACGTTCACCACCAT | qPCR of the *csuA/B* gene | |
| *3773*_qPCR_FW | AGTGTCACTGCGGGTTACTG | qPCR of the *ABUW_3773* gene | |
| *3773*_qPCR_RV | CTAGGTTGTCCCGCCTCATC | qPCR of the *ABUW_3773* gene | |
| Hyg_FW | TCATCACTGCAGATGAAAAAGCCTGAACTCACC | Cloning of *hph* cassette in mini-Tn7T. | |
| Hyg_RV | TCATCACTGCAGGGGGGATCGATCCCGGTCGG | Cloning of *hph* cassette in mini-Tn7T. | |
| *abaM*_compl_FW_*Bam*HI | ATATGGATCCTTGCTCTCATTAGACTCCATTCAC | *abaM*::T26 complementation. | |
| *abaM*_compl_RV_*Kpn*I | ATATGGTACCGTGCTTCCACTTATTTTTCAAGT | *abaM*::T26 complementation. | |
| *abaR*_RV | CTACAAAAGCCCTAGCATTACAGC | RT-PCRs (*abaR*-*abaM*) | |
| *abaM*_FW | GGTTAGCATACCCCCTCATTTC | RT-PCRs (*abaM* and *abaM*-*abaI*) | |
| *abaM*_RV | GGTTTGACTTAATGAAGACTCG | RT-PCRs (*abaM* and *abaR*-*abaM*) | |
| *abaI*_FW | CAATTTTTCAGAAGGCCTATATACC | RT-PCRs (*abaI)* | |
| *abaI*_RV | CAATCAAGCATGCAAACATC | RT-PCRs (*abaI* and *abaM*-*abaI*) | |

**Table S3:** Differentially expressed genes (log_2_(fold change) ≥ 1) in *abaI*::T26 (compared with AB5075 wild-type).

| Gene id | Log_2_(Fold change) | Description | NCBI Protein Accession |
| --- | --- | --- | --- |
| *ABUW_RS18385* | 4.3142 | *N*-acylhomoserine lactone synthase (*abaI*) | WP_001020940.1 |
| *ABUW_RS07260* | 4.2853 | SCPU domain-containing protein (*csuB*) | WP_000876475.1 |
| *ABUW_RS07255* | 4.1072 | protein CsuA (*csuA*) | WP_000577009.1 |
| *ABUW_RS07265* | 3.9262 | molecular chaperone (*csuC*) | WP_001065473.1 |
| *ABUW_RS07250* | 3.9062 | SCPU domain-containing protein (*csuA/B*) | WP_000790104.1 |
| *ABUW_RS07275* | 3.692 | protein CsuE (*csuE*) | WP_002017500.1 |
| *ABUW_RS07270* | 3.6287 | fimbrial biogenesis outer membrane usher protein (*csuD*) | WP_000603294.1 |
| *ABUW_RS07245* | 2.8129 | TetR/AcrR family transcriptional regulator | WP_000590096.1 |
| *ABUW_RS07925* | 1.9432 | alcohol dehydrogenase | WP_000874704.1 |
| *ABUW_RS01970* | 1.9355 | flavohemoprotein | WP_000188888.1 |
| *ABUW_RS07910* | 1.9263 | aldehyde dehydrogenase | WP_001269058.1 |
| *ABUW_RS07630* | 1.8189 | hypothetical protein | WP_000783724.1 |
| *ABUW_RS07640* | 1.7361 | hypothetical protein | WP_000772607.1 |
| *ABUW_RS07620* | 1.5361 | hypothetical protein | WP_000020771.1 |
| *ABUW_RS07905* | 1.5245 | ethanolamine permease | WP_001075439.1 |
| *ABUW_RS07625* | 1.4155 | LLM class flavin-dependent oxidoreductase | WP_001257199.1 |
| *ABUW_RS07240* | 1.4129 | hypothetical protein | WP_000637306.1 |
| *ABUW_RS08275* | 1.402 | DNA methylase | WP_000652017.1 |
| *ABUW_RS06205* | 1.3599 | DNA helicase | WP_000106166.1 |
| *ABUW_RS07280* | 1.3354 | hypothetical protein | WP_000821199.1 |
| *ABUW_RS08025* | 1.3233 | hypothetical protein | WP_001092426.1 |
| *ABUW_RS07895* | 1.3225 | ethanolamine ammonia-lyase subunit EutC | WP_000774022.1 |
| *ABUW_RS19740* | 1.3116 | hypothetical protein | WP_002017532.1 |
| *ABUW_RS08010* | 1.3043 | hypothetical protein | WP_001243508.1 |
| *ABUW_RS07980* | 1.2832 | glutathione S-transferase family protein | WP_001279985.1 |
| *ABUW_RS07800* | 1.2821 | membrane protein | WP_001228338.1 |
| *ABUW_RS08160* | 1.2796 | 3-oxoacyl-ACP reductase | WP_000132115.1 |
| *ABUW_RS07445* | 1.2788 | dicarboxylate/amino acid:cation symporter | WP_000347180.1 |
| *ABUW_RS08155* | 1.2734 | acetyl-CoA C-acyltransferase | WP_000047940.1 |
| *ABUW_RS05005* | 1.2604 | alpha/beta hydrolase | WP_000149921.1 |
| *ABUW_RS08165* | 1.2485 | hypothetical protein | WP_001097352.1 |
| *ABUW_RS08145* | 1.2425 | LysR family transcriptional regulator | WP_001024733.1 |
| *ABUW_RS19150* | 1.2359 | hypothetical protein | WP_001024502.1 |
| *ABUW_RS11580* | 1.2324 | taurine ABC transporter permease | WP_001092656.1 |
| *ABUW_RS07985* | 1.2211 | short-chain dehydrogenase | WP_000135428.1 |
| *ABUW_RS08190* | 1.2166 | phospholipase C%2C phosphocholine-specific | WP_000633003.1 |
| *ABUW_RS07715* | 1.2132 | 2-amino-4-hydroxy-6-hydroxymethyldihydropteridine pyrophosphokinase | WP_000993407.1 |
| *ABUW_RS08170* | 1.2116 | esterase | WP_000588779.1 |
| *ABUW_RS19155* | 1.2107 | hypothetical protein | WP_000241707.1 |
| *ABUW_RS07725* | 1.2067 | K(+)-transporting ATPase subunit B | WP_001097170.1 |
| *ABUW_RS07435* | 1.186 | MFS transporter | WP_001066110.1 |
| *ABUW_RS11590* | 1.1859 | taurine ABC transporter substrate-binding protein | WP_000105187.1 |
| *ABUW_RS19145* | 1.1712 | transcriptional regulator | WP_000096600.1 |
| *ABUW_RS07920* | 1.1707 | alpha/beta hydrolase | WP_001098135.1 |
| *ABUW_RS07955* | 1.1695 | aminopeptidase N | WP_001017234.1 |
| *ABUW_RS05010* | 1.1572 | sulfate ABC transporter substrate-binding protein | WP_001212600.1 |
| *ABUW_RS07845* | 1.1519 | tRNA (adenosine(37)-N6)-threonylcarbamoyltransferase complex ATPase subunit type 1 TsaE | WP_031944287.1 |
| *ABUW_RS08125* | 1.1426 | membrane protein | WP_001169538.1 |
| *ABUW_RS07465* | 1.1379 | methionine ABC transporter ATP-binding protein | WP_000614025.1 |
| *ABUW_RS19770* | 1.1377 | hypothetical protein | - |
| *ABUW_RS08255* | 1.132 | hypothetical protein | WP_001238662.1 |
| *ABUW_RS19165* | 1.1273 | toxic anion resistance protein | WP_000933769.1 |
| *ABUW_RS07360* | 1.123 | acyl-CoA desaturase | WP_000032710.1 |
| *ABUW_RS19170* | 1.1197 | hypothetical protein | WP_001093570.1 |
| *ABUW_RS07935* | 1.1175 | 5-carboxymethyl-2-hydroxymuconate isomerase | WP_001120984.1 |
| *ABUW_RS07460* | 1.1162 | ABC transporter permease | WP_001205190.1 |
| *ABUW_RS07505* | 1.1138 | tRNA uridine-5-carboxymethylaminomethyl(34) synthesis enzyme MnmG | WP_000559187.1 |
| *ABUW_RS08225* | 1.1107 | alpha/beta hydrolase | WP_001290021.1 |
| *ABUW_RS19105* | 1.1103 | hypothetical protein | WP_001070890.1 |
| *ABUW_RS19075* | 1.1051 | hypothetical protein | WP_001012847.1 |
| *ABUW_RS19080* | 1.1038 | thermonuclease | WP_000861205.1 |
| *ABUW_RS07735* | 1.1031 | sensor histidine kinase KdpD | WP_001191611.1 |
| *ABUW_RS07660* | 1.0966 | peptide chain release factor N(5)-glutamine methyltransferase | WP_001017509.1 |
| *ABUW_RS07605* | 1.0929 | poly-beta-1%2C6-N-acetyl-D-glucosamine N-deacetylase PgaB | WP_001061302.1 |
| *ABUW_RS07775* | 1.0919 | hypothetical protein | WP_000756505.1 |
| *ABUW_RS07650* | 1.0916 | type 1 glutamine amidotransferase domain-containing protein | WP_000735895.1 |
| *ABUW_RS07395* | 1.0911 | DNA-binding response regulator | WP_000526534.1 |
| *ABUW_RS08035* | 1.0891 | TonB-dependent siderophore receptor | WP_000527015.1 |
| *ABUW_RS07720* | 1.0799 | potassium-transporting ATPase subunit KdpA | WP_000891202.1 |
| *ABUW_RS07305* | 1.0773 | TetR/AcrR family transcriptional regulator | WP_000792929.1 |
| *ABUW_RS19160* | 1.0754 | hypothetical protein | WP_000661588.1 |
| *ABUW_RS08095* | 1.0722 | universal stress protein | WP_001109442.1 |
| *ABUW_RS07785* | 1.0593 | hypothetical protein | WP_000920881.1 |
| *ABUW_RS08250* | 1.0587 | MFS transporter | WP_071543520.1 |
| *ABUW_RS08235* | 1.0544 | PepSY domain-containing protein | WP_000075461.1 |
| *ABUW_RS08280* | 1.0528 | nitrate transporter | WP_000039924.1 |
| *ABUW_RS07235* | 1.052 | FadR family transcriptional regulator | WP_000572518.1 |
| *ABUW_RS07840* | 1.05 | pseudouridylate synthase | WP_000097958.1 |
| *ABUW_RS07990* | 1.0454 | oxidoreductase | WP_000920714.1 |
| *ABUW_RS08110* | 1.0389 | RDD family protein | WP_000221476.1 |
| *ABUW_RS07710* | 1.0383 | dihydroneopterin aldolase | WP_000338779.1 |
| *ABUW_RS07600* | 1.0353 | poly-beta-1%2C6 N-acetyl-D-glucosamine export porin PgaA | WP_000913301.1 |
| *ABUW_RS07560* | 1.035 | protoheme IX farnesyltransferase | WP_000915319.1 |
| *ABUW_RS19300* | 1.0227 | aminoglycoside O-phosphotransferase APH(3'')-Ib | WP_025464697.1 |
| *ABUW_RS07930* | 1.0224 | helicase | WP_000808297.1 |
| *ABUW_RS08090* | 1.0174 | ABC transporter ATP-binding protein | WP_000193596.1 |
| *ABUW_RS08185* | 1.0159 | DNA polymerase III subunit gamma/tau | WP_045887671.1 |
| *ABUW_RS07815* | 0.99981 | DUF441 domain-containing protein | WP_000880863.1 |
| *ABUW_RS15330* | -1.016 | hypothetical protein | WP_001001666.1 |
| *ABUW_RS09690* | -1.0661 | zonular occludens toxin | WP_032017212.1 |
| *ABUW_RS13220* | -1.0776 | hypothetical protein | WP_001037898.1 |
| *ABUW_RS15280* | -1.1274 | hypothetical protein | WP_000206132.1 |
| *ABUW_RS09695* | -1.1693 | hypothetical protein | WP_032017213.1 |
| *ABUW_RS01270* | -1.1982 | sulfate permease | WP_001111063.1 |
| *ABUW_RS09700* | -1.2079 | hypothetical protein | WP_032017215.1 |
| *ABUW_RS01160* | -1.2284 | aquaporin Z | WP_001045986.1 |
| *ABUW_RS12990* | -1.4531 | hypothetical protein | WP_001034729.1 |

**Table S4**: Differentially expressed genes (log_2_(fold change) ≥ 1) in *abaM*::T26 (compared with AB5075 wild-type)

| Gene id | Log_2_(Fold change) | Description | NCBI Protein Accession |
| --- | --- | --- | --- |
| *ABUW_RS07245* | 2.9052 | TetR/AcrR family transcriptional regulator | WP_000590096.1 |
| *ABUW_RS07260* | 2.878 | SCPU domain-containing protein (*csuB*) | WP_000876475.1 |
| *ABUW_RS18365* | 2.8531 | acyl-CoA dehydrogenase (*ABUW_3772*) Ac-505 biosynthetic operon | WP_000060267.1 |
| *ABUW_RS11870* | 2.7441 | hypothetical protein | WP_000108365.1 |
| *ABUW_RS18350* | 2.6991 | RND transporter (*ABUW_3769*).  Ac-505 biosynthetic operon | WP_000256252.1 |
| *ABUW_RS18355* | 2.6293 | non-ribosomal peptide synthetase (ABUW_3770) | WP_001060991.1 |
| *ABUW_RS18375* | 2.554 | LuxR family transcriptional regulator (*abaR*) | WP_000446790.1 |
| *ABUW_RS07265* | 2.5191 | molecular chaperone (*csuC*) | WP_001065473.1 |
| *ABUW_RS07250* | 2.4684 | SCPU domain-containing protein (*csuA/B*) | WP_000790104.1 |
| *ABUW_RS18385* | 2.4087 | N-acylhomoserine lactone synthase (*abaI*) | WP_001020940.1 |
| *ABUW_RS18370* | 2.3341 | acyl-CoA synthetase (*ABUW_3773*) Ac-505 biosynthetic operon | WP_000279948.1 |
| *ABUW_RS07255* | 2.323 | protein CsuA (*csuA*) | WP_000577009.1 |
| *ABUW_RS07270* | 2.2652 | fimbrial biogenesis outer membrane usher protein (*csuD*) | WP_000603294.1 |
| *ABUW_RS07275* | 2.1508 | protein CsuE (*csuE*) | WP_002017500.1 |
| *ABUW_RS07145* | 1.8615 | DUF2171 domain-containing protein | WP_001094391.1 |
| *ABUW_RS11840* | 1.7989 | stress-induced protein | WP_000024222.1 |
| *ABUW_RS08050* | 1.7574 | hypothetical protein | WP_001123841.1 |
| *ABUW_RS11875* | 1.7165 | hypothetical protein | WP_000132046.1 |
| *ABUW_RS19150* | 1.5586 | hypothetical protein | WP_001024502.1 |
| *ABUW_RS19060* | 1.4374 | hypothetical protein | WP_000910917.1 |
| *ABUW_RS19165* | 1.4373 | toxic anion resistance protein | WP_000933769.1 |
| *ABUW_RS19145* | 1.4181 | transcriptional regulator | WP_000096600.1 |
| *ABUW_RS03420* | 1.4179 | tRNA-Asp | - |
| *ABUW_RS01970* | 1.4088 | flavohemoprotein | WP_000188888.1 |
| *ABUW_RS19155* | 1.3734 | hypothetical protein | WP_000241707.1 |
| *ABUW_RS19115* | 1.3711 | hypothetical protein | WP_001057663.1 |
| *ABUW_RS19170* | 1.3708 | hypothetical protein | WP_001093570.1 |
| *ABUW_RS19140* | 1.3556 | ParA family protein | WP_000724905.1 |
| *ABUW_RS13115* | 1.3406 | hypothetical protein | WP_000980495.1 |
| *ABUW_RS19080* | 1.2162 | thermonuclease | WP_000861205.1 |
| *ABUW_RS19240* | 1.202 | hypothetical protein | - |
| *ABUW_RS19305* | 1.1854 | ANT(3'')-Ia family aminoglycoside nucleotidyltransferase AadA2 | WP_001261740.1 |
| *ABUW_RS20015* | 1.1825 | hypothetical protein | - |
| *ABUW_RS19065* | 1.1742 | hypothetical protein | WP_000095825.1 |
| *ABUW_RS12220* | 1.1605 | hypothetical protein | WP_000008105.1 |
| *ABUW_RS19105* | 1.1574 | hypothetical protein | WP_001070890.1 |
| *ABUW_RS11885* | 1.1476 | hypothetical protein | WP_001136759.1 |
| *ABUW_RS13925* | 1.1447 | 23S rRNA (pseudouridine(1915)-N(3))-methyltransferase RlmH | WP_000702193.1 |
| *ABUW_RS20020* | 1.1374 | transposition protein TniB | WP_001381192.1 |
| *ABUW_RS19595* | 1.0943 | DNA-binding protein | WP_001096616.1 |
| *ABUW_RS19110* | 1.0805 | DUF2786 domain-containing protein | WP_000389915.1 |
| *ABUW_RS19200* | 1.0625 | hypothetical protein | WP_000790084.1 |
| *ABUW_RS19820* | 1.0579 | hypothetical protein | - |
| *ABUW_RS19190* | 1.0498 | hypothetical protein | WP_000701003.1 |
| *ABUW_RS19235* | 1.0446 | hypothetical protein | WP_000654348.1 |
| *ABUW_RS19180* | 1.0398 | hypothetical protein | WP_000443897.1 |
| *ABUW_RS10070* | 1.0279 | Fur family transcriptional regulator | WP_000207886.1 |
| *ABUW_RS13010* | 1.0165 | DUF4142 domain-containing protein | WP_000644336.1 |
| *ABUW_RS19345* | 1.0122 | molecular chaperone DnaJ | WP_000758689.1 |
| *ABUW_RS08580* | 1.009 | DUF159 family protein | WP_000332547.1 |
| *ABUW_RS19300* | 1.0071 | aminoglycoside O-phosphotransferase APH(3'')-Ib | WP_025464697.1 |
| *ABUW_RS11975* | 1.003 | PaaI family thioesterase | WP_000445634.1 |
| *ABUW_RS10685* | -0.99885 | HU family DNA-binding protein | WP_001043034.1 |
| *ABUW_RS14775* | -1.0208 | outer membrane protein | WP_001202415.1 |
| *ABUW_RS17210* | -1.0241 | Na+/H+ antiporter subunit E | WP_001177202.1 |
| *ABUW_RS18725* | -1.045 | thiol:disulfide interchange protein DsbA/DsbL | WP_000737570.1 |
| *ABUW_RS15925* | -1.0539 | integration host factor subunit alpha | WP_000126166.1 |
| *ABUW_RS03825* | -1.0607 | hypothetical protein | WP_001278744.1 |
| *ABUW_RS15280* | -1.0657 | hypothetical protein | WP_000206132.1 |
| *ABUW_RS18190* | -1.0826 | ATP synthase subunit C | WP_000424060.1 |
| *ABUW_RS08725* | -1.0884 | hypothetical protein | WP_000179560.1 |
| *ABUW_RS02765* | -1.0913 | carbapenem-hydrolyzing class D beta-lactamase OXA-23 | WP_001046004.1 |
| *ABUW_RS03465* | -1.0935 | alanine:cation symporter family protein | WP_001005337.1 |
| *ABUW_RS09695* | -1.0973 | hypothetical protein | WP_032017213.1 |
| *ABUW_RS17205* | -1.1655 | monovalent cation/H+ antiporter subunit D | WP_000459267.1 |
| *ABUW_RS04295* | -1.1868 | succinyl-CoA ligase subunit beta | WP_001048573.1 |
| *ABUW_RS01270* | -1.2922 | sulfate permease | WP_001111063.1 |
| *ABUW_RS13140* | -1.3221 | heavy metal-associated domain protein | WP_000770719.1 |
| *ABUW_RS14740* | -1.3499 | hemerythrin | WP_000782976.1 |
| *ABUW_RS15385* | -1.35 | NADH-quinone oxidoreductase subunit K | WP_000529822.1 |
| *ABUW_RS17440* | -1.4553 | membrane protein | WP_000472947.1 |
| *ABUW_RS03380* | -1.5003 | entericidin%2C EcnA/B family | WP_000757214.1 |
| *ABUW_RS17200* | -1.5374 | Na+/H+ antiporter subunit C | WP_000624229.1 |
| *ABUW_RS02775* | -1.8507 | phage tail assembly protein | WP_001071615.1 |
| *ABUW_RS12990* | -2.1063 | hypothetical protein | WP_001034729.1 |
| *ABUW_RS00585* | -2.1307 | hypothetical protein | WP_000770763.1 |

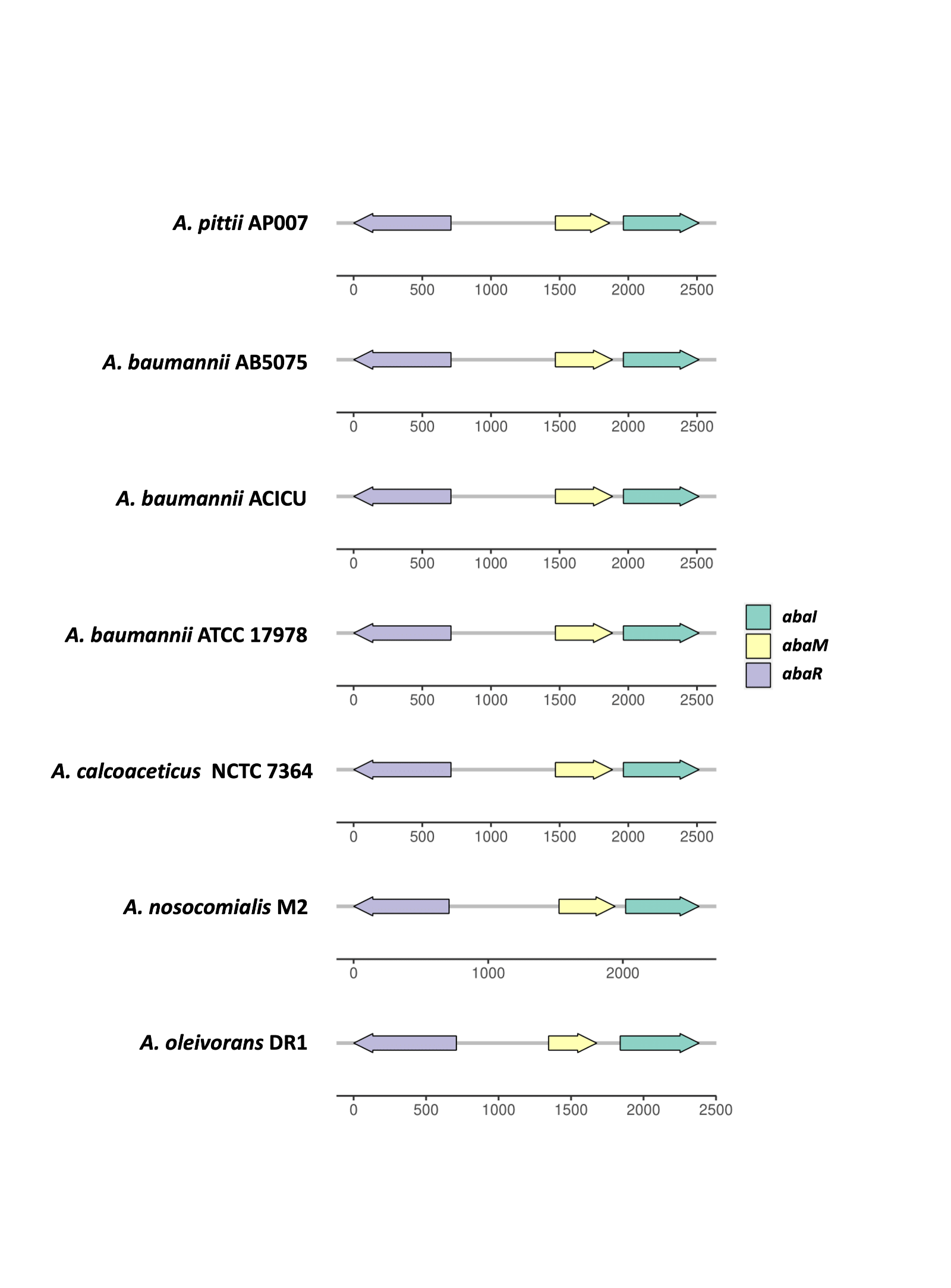

FIG S1 Conservation of quorum sensing locus organization among different *Acinetobacter spp.*

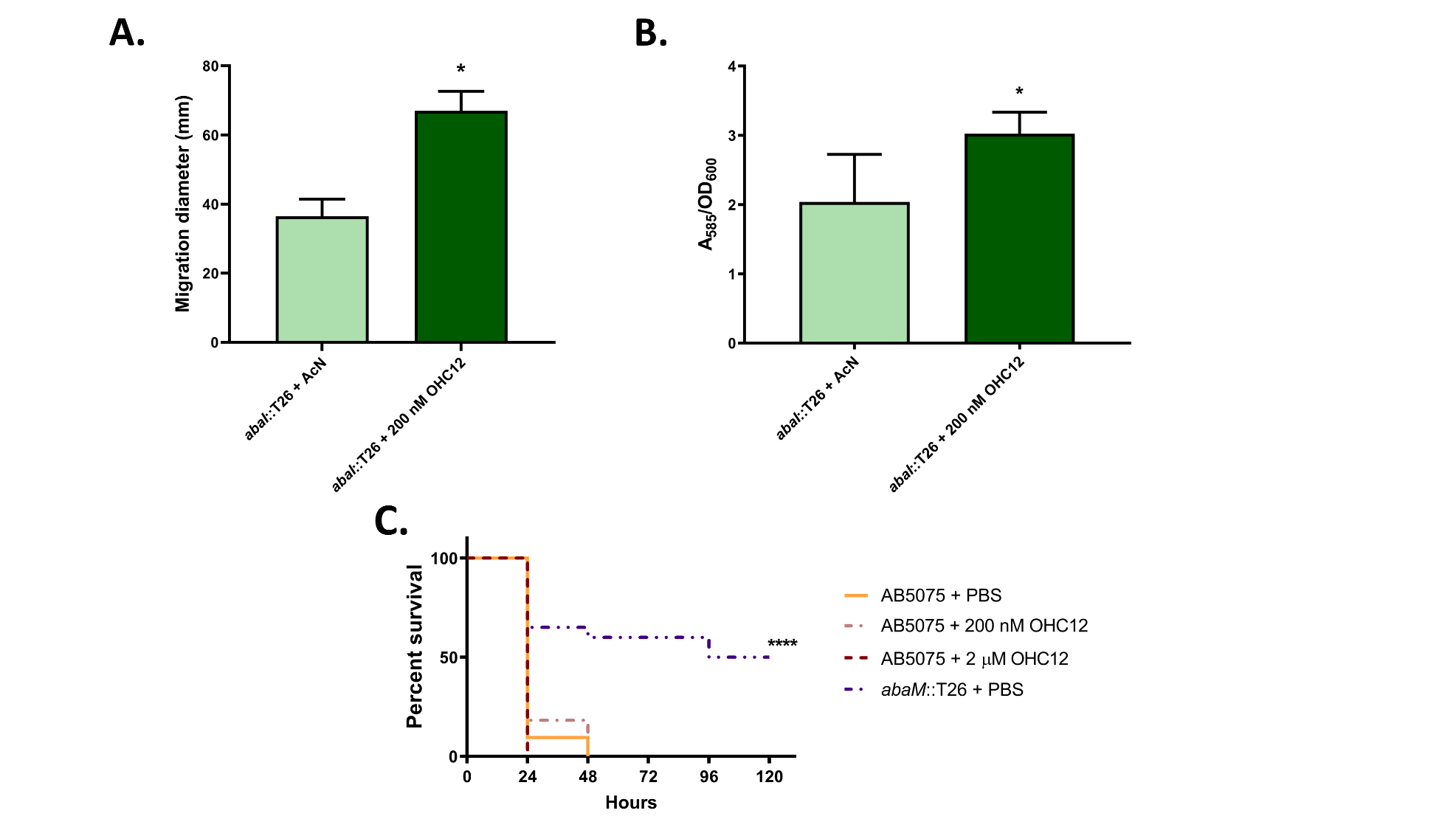

**FIG S2** Contribution of OHC12 to AB5075 phenotypes. **(A)** Surface motility and (**B**) biofilm formation on polypropylene by the *abaI* mutant with and without OHC12. Acn, Acetonitrile solvent control. **(C)** *Galleria mellonella* larvae killing by wild type with and without OHC12 compared with the *abaM* mutant after inoculation of approximately 2 x 10^4^ CFU/larvae. Asterisks indicate statistically significant differences compared to the wild-type AB5075 strain: **, p ≤ 0.01; ***, p ≤ 0.001; ****; p ≤ 0.0001

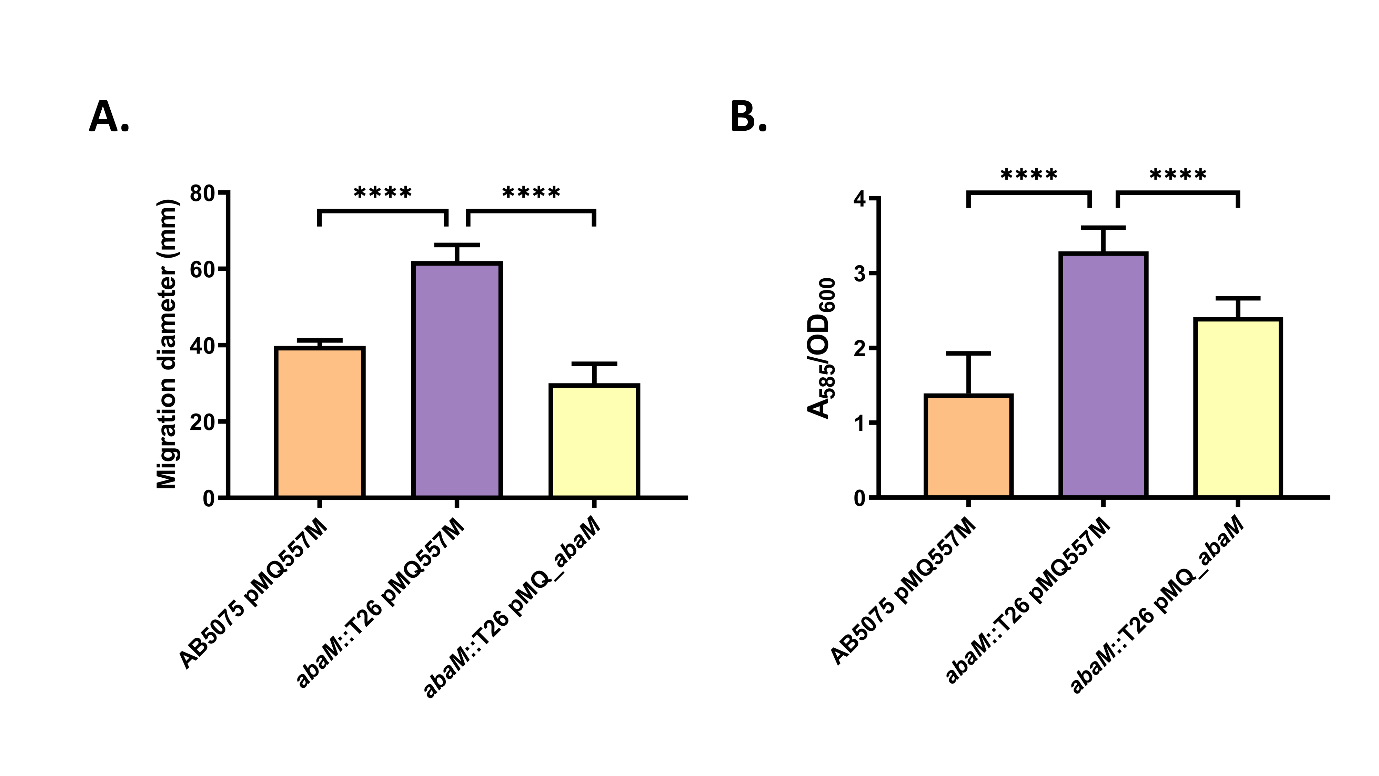

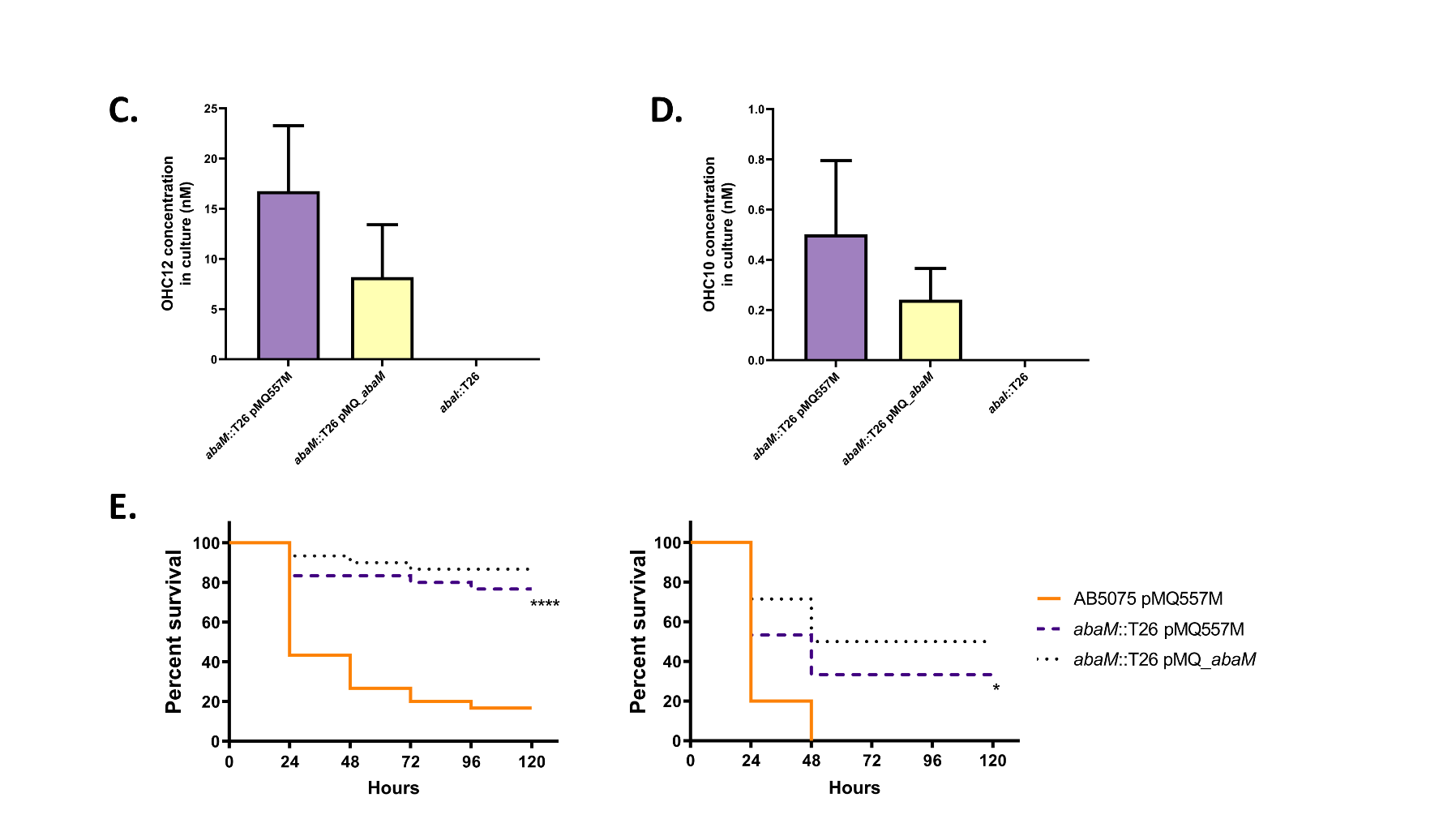

FIG S3 Genetic complementation of the *abaM*::T26 mutant. (A) Surface motility in 0.3% Eiken agar LS-LB plates. (B) Biofilm formation in polypropylene. (C) OHC12 production.
(D) OHC10 production. (E) *Galleria mellonella* larvae killing after inoculation of approximately 2 x 10^4^ (left) or 2 x 10^5^ (right) CFU/larva. Asterisks indicate statistically significant differences compared with the wild-type AB5075 strain: *, p ≤ 0.05; ****, p ≤ 0.0001.

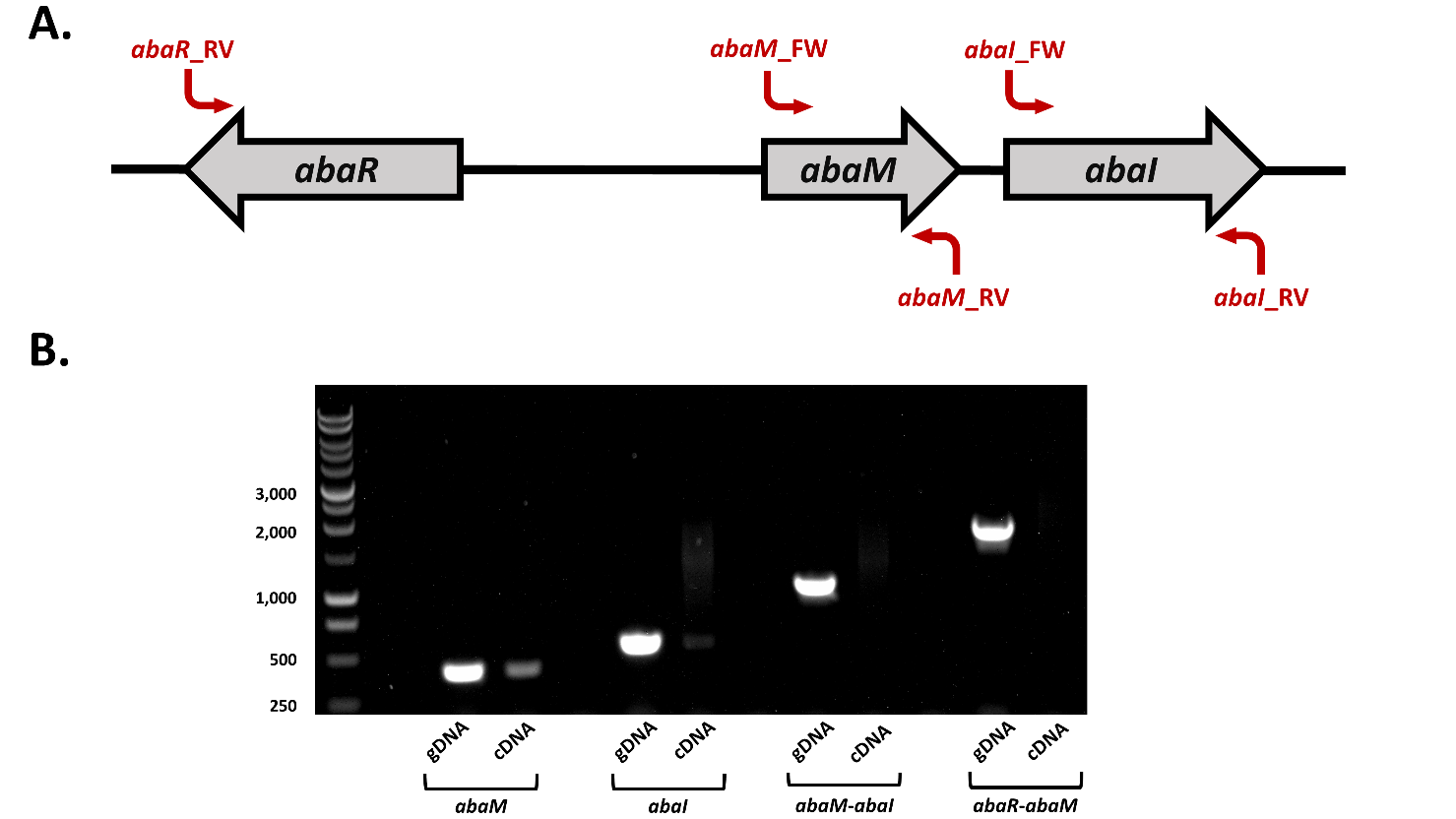

**Fig. S4** The *abaM* and *abaI* genes are not co-transcribed. **(A)** Organization of the *abaRMI* region and the positions of the PCR primers used **(B)** Agarose gel showing single *abaM* and *abaI* transcripts. No cDNAs were obtained for *abaM-abaI* or for *abaR*-*abaM*. The left hand lane shows the DNA molecular markers. gDNA, genomic DNA control. The *abaM*_FW and *abaM*_RV primer pairs were used to amplify the *abaM* gene; *abaI*_FW and *abaI*_RV pair for the *abaI* gene, the *abaM*_FW and *abaI*_RV pair for the *abaM*-*abaI* region and the *abaR*_RV and *abaM*_RV pair for *abaR*-*abaM*.

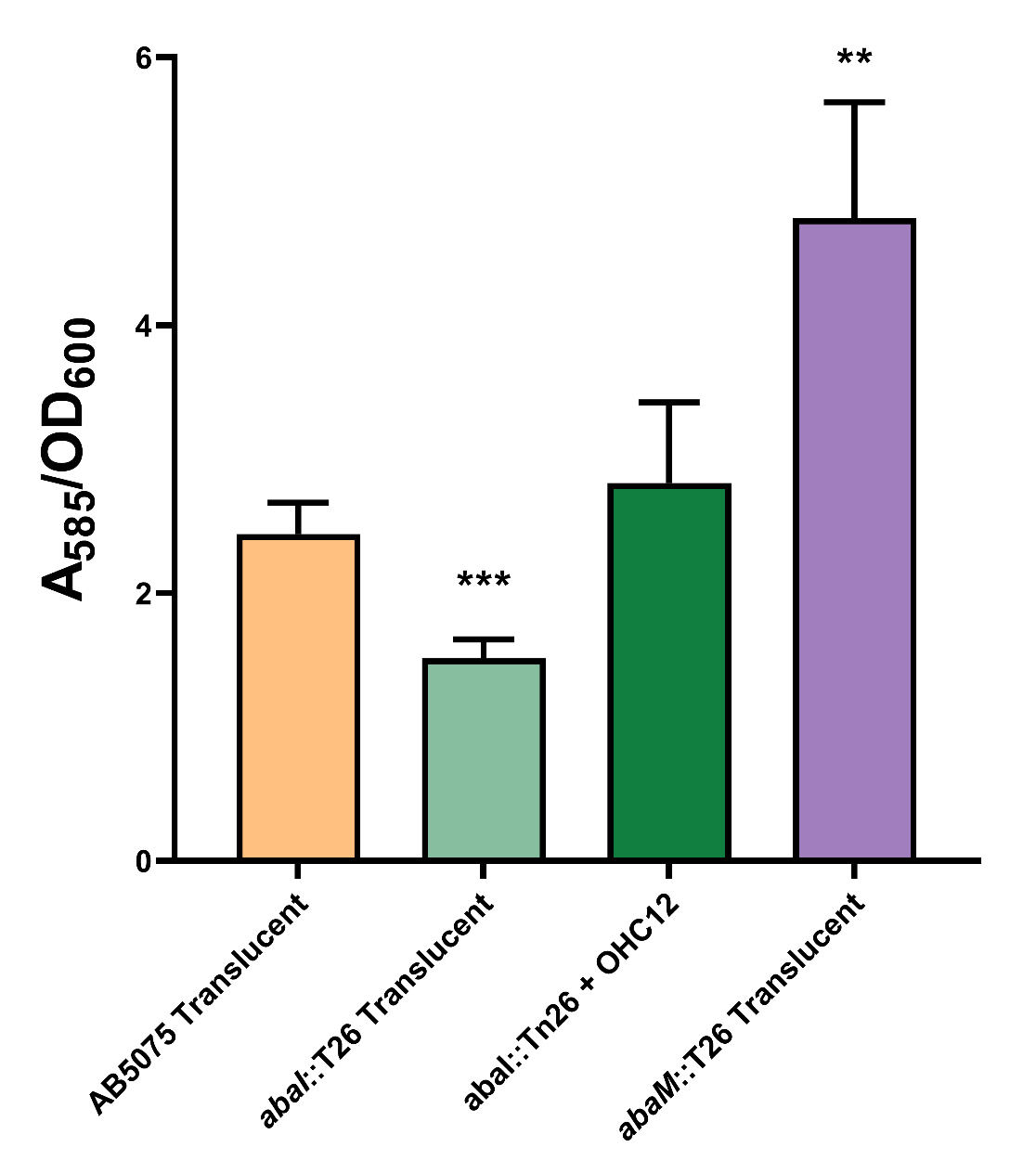

FIG S5 Biofilm formation on polypropylene tubes by translucent variants of AB5075 wild-type, *abaI*::T26, and *abaM*::T26. The *abaI*::T26 mutant was also supplemented with OHC12 (200 nM). Asterisks indicate statistically significant differences compared to the wild-type AB5075 strain: **, p ≤ 0.01; ****; p ≤ 0.0001.

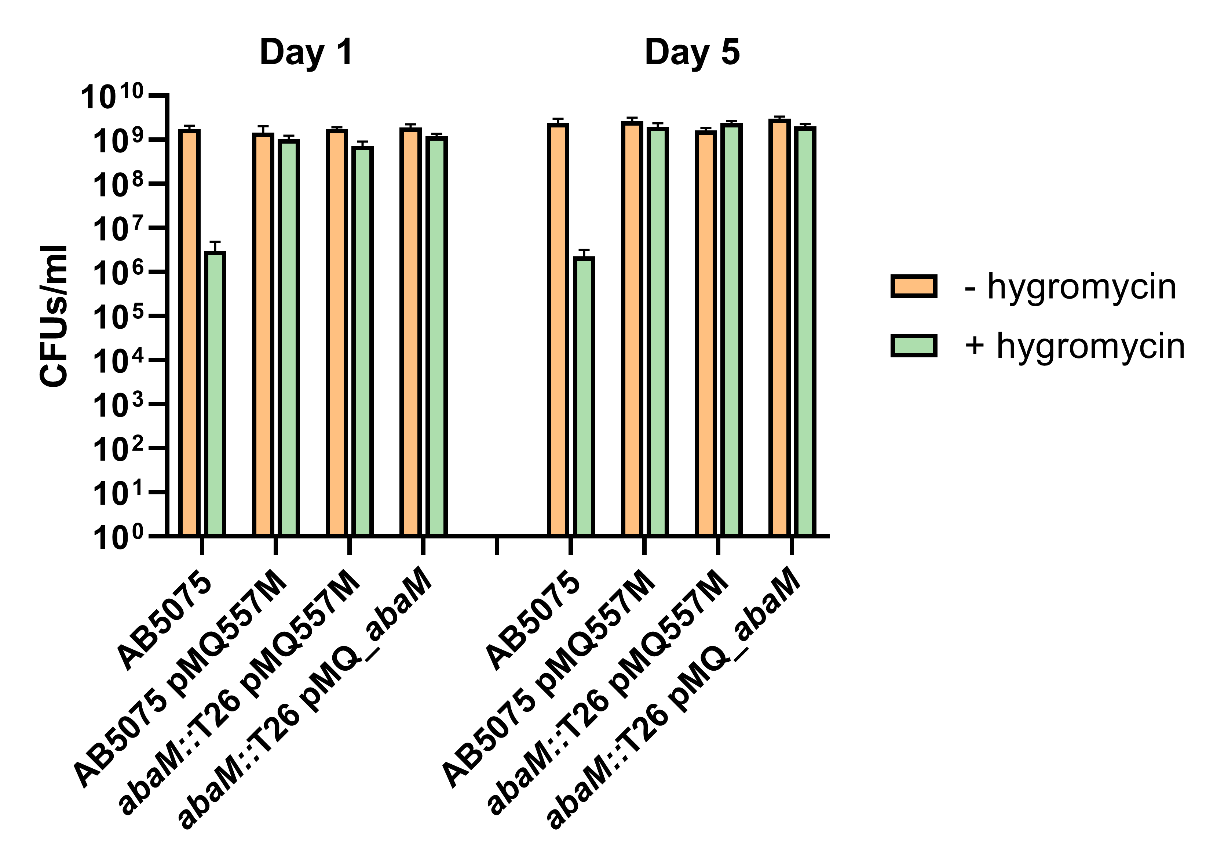

**FIG S6** Stability of the vector pMQ557M and *abaM* complementing plasmid pMQabaM in the wild type AB5075 and *abaM* mutant in the presence and absence of hygromycin selection after 5 days of repeat daily subculturing.
